# TGF-β signaling is critical for maintenance of the tendon cell fate

**DOI:** 10.1101/823021

**Authors:** Guak-Kim Tan, Brian A. Pryce, Anna Stabio, John V. Brigande, ChaoJie Wang, Zheng Xia, Sara F. Tufa, Douglas R. Keene, Ronen Schweitzer

## Abstract

Studies of cell fate focus on specification, but little is known about maintenance of the differentiated state. We find that TGFβ signaling plays an essential role in maintenance of the tendon cell fate. To examine the role TGFβ signaling in tenocytes TGFβ type II receptor was targeted in the Scleraxis cell lineage. Tendon development was not disrupted in mutant embryos, but shortly after birth tenocytes lost differentiation markers and reverted to a more stem/progenitor state. Targeting of Tgfbr2 using other Cre drivers did not cause tenocyte dedifferentiation suggesting a critical significance for the spatio-temporal activity of ScxCre. Viral reintroduction of Tgfbr2 to mutants was sufficient to prevent and even rescue mutant tenocytes suggesting a continuous and cell-autonomous role for TGFβ signaling in cell fate maintenance. These results uncover the critical importance of molecular pathways that maintain the differentiated cell fate and a key role for TGFβ signaling in these processes.

## Introduction

Studies of cell fate determination are in most cases focused on the signaling pathways and transcription factors that direct naïve cells to assume a specific cell fate (Li et al., 2012; James, 2013; Huang et al., 2015). It is commonly accepted that once fully differentiated the cells enter a stable cellular phenotype, but relatively little is known about the molecular mechanisms that reinforce and maintain this differentiated state. Maintenance of the differentiated state is however essential for tissue function and identifying the molecular pathways involved in these processes may be of great importance for understanding tissue homeostasis and pathology.

Tendons are connective tissues that transmit forces from muscle to bone to generate movement (Kannus, 2000). Despite their importance to overall musculoskeletal function and their slow and limited healing capabilities, relatively little is known about tendon development, the tendon cell fate, maturation and pathology. Elucidating the key molecular regulators of these processes is thus essential for improvements in the management of tendon healing, the treatment of tendinopathy and for bioengineering efforts for this tissue.

A limited number of transcription factors were so far identified as key regulators of the tendon cell fate including most notably, Scleraxis (Scx), a bHLH transcription factor expressed in tendon cells from progenitor stages and through development (Schweitzer et al., 2001) and Mohawk (Mkx), an atypical homeobox protein with essential roles in the development of the collagen matrix in tendons (Ito et al., 2010). Prototypic markers for the tendon cell fate also include the transmembrane protein tenomodulin (Tnmd) and collagen type I (Kannus, 2000; Huang et al., 2015), the major building blocks of the tendon fibrillar extracellular matrix that mediates the transmission of force by tendons.

Previous studies have also established a central role for the transforming growth factor-β (TGFβ) signaling pathway in early events of tendon development (Pryce et al., 2009; Havis et al., 2016). Notably, TGFβ is a potent inducer of Scleraxis (Scx) both *in vivo* and in cultured cells and disruption of TGFβ signaling in mouse limb bud mesenchyme resulted in complete failure of tendon formation (Pryce et al., 2009). This phenotype manifested at the onset of embryonic tendon development but robust expression of TGFβ ligands and associated molecules in later stages of tendon development suggested possible additional roles for TGFβ signaling in tendon development (Kuo et al., 2008; Pryce et al., 2009). Moreover, subcutaneous application of growth and differentiation factors (GDFs), members of the TGFβ superfamily, can induce ectopic neo-tendon formation in rats (Wolfman et al., 1997). The goal of this study was therefore to ask if TGFβ signaling plays essential roles at later stages of tendon development.

The TGFβ superfamily comprises secreted polypeptides that regulate diverse developmental processes ranging from cellular growth, differentiation and migration to tissue patterning and morphogenesis (Santibanez et al., 2011; Sakaki-Yumoto et al., 2013). These ligands act by binding to transmembrane type II receptors, which in turn recruit and activate a type I receptor. The activated receptor complex subsequently phosphorylates and activates receptor-regulated transcription factors called Smads (Smad2/3 for TGFβ signaling) that subsequently complex with the common-mediator Smad4 and translocate into the nucleus where they promote or repress responsive target genes (Vander Ark et al., 2018). The TGFβ proper ligands (TGFβ1-3) all bind to a single type II receptor. Consequently, disrupting this one receptor is sufficient to abrogate all TGFβ signaling. To test for additional roles of TGFβ signaling in tendon development and biology we wanted to bypass the early essential function in tendon formation, and decided to target TGFβ type II receptor (Tgfbr2) explicitly in tendon cells. We therefore targeted the receptor using ScxCre (Blitz et al., 2013), a tendon specific Cre driver, so that TGFβ signaling will be disrupted specifically in tendon cells and only after the initial events of tendon formation.

We find that tendon differentiation function and growth during embryonic development was not disrupted following targeted deletion of TGFβ signaling in tenocytes, but shortly after birth the cells lost tendon cell differentiation markers and reverted to a more progenitor-like state. Moreover, viral reintroduction of Tgfbr2 to mutant cells was sufficient to prevent dedifferentiation and even to rescue the tendon cell fate in a cell-autonomous manner, highlighting a continuous and essential role of TGFβ signaling in maintenance of the tendon cell fate.

## Results

### Targeting TGFβ type II receptor in Scx-expressing cells resulted in tendon disruption and limb abduction

Our previous studies showed that disruption of TGFβ signaling in limb mesenchyme resulted in the complete failure of tendon formation (Pryce et al., 2009). To examine later roles of TGFβ signaling in tendon development we targeted Tgfbr2 with ScxCre thereby bypassing the early role of TGFβ signaling for tendon development. Tgfbr2;ScxCre mutant embryos indeed developed a complete network of tendons by E14.5, indicating they have bypassed the early requirement for TGFβ signaling in tendon development (Fig. 1A).

**Fig 1.**
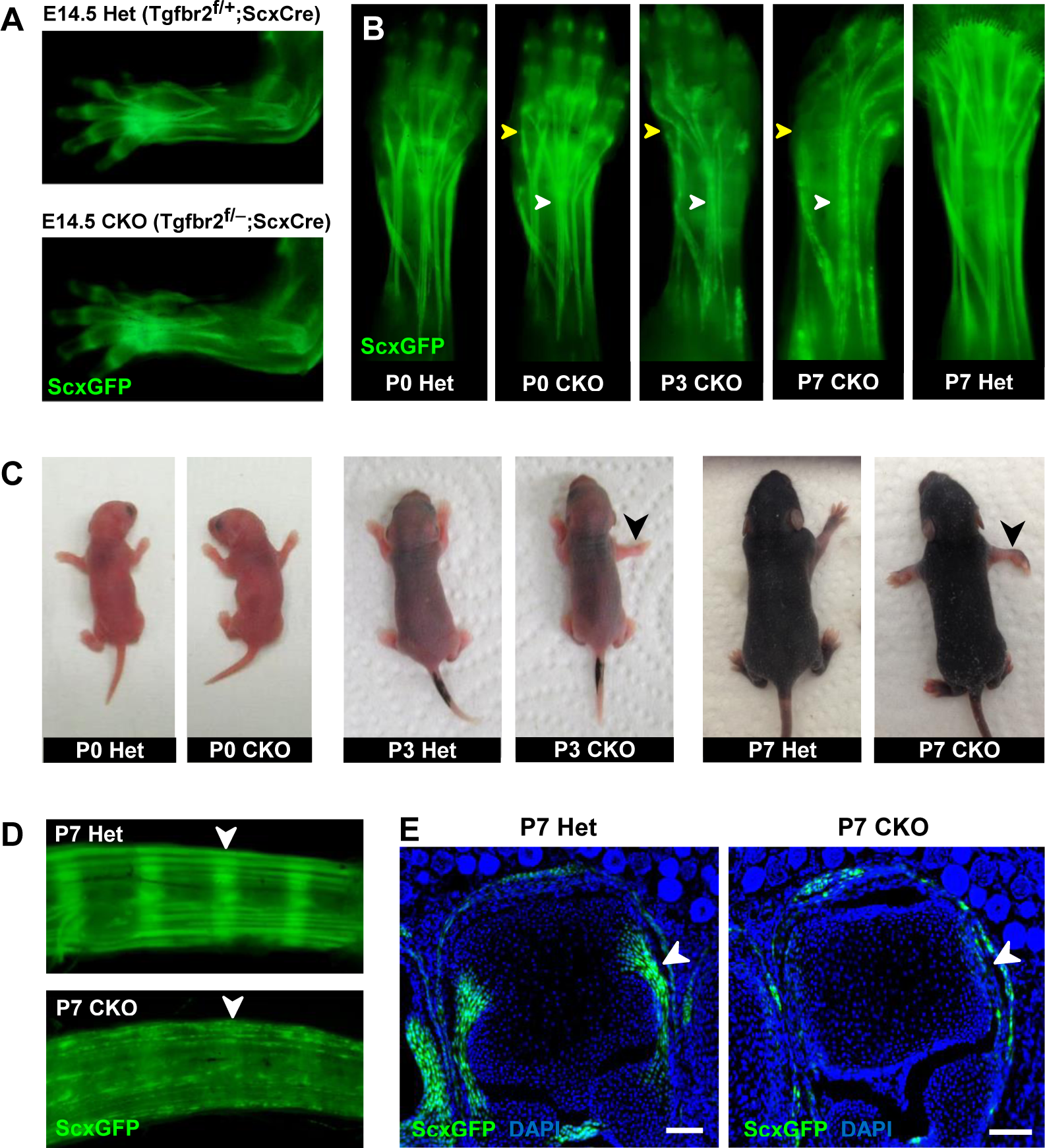
Tendon phenotypes manifested in Tgfbr2;ScxCre mutants. (A-D) Whole-mount imaging in fluorescent ScxGFP signal or brightfield. (A) Dorsally-viewed embryo forelimb shows the formation of a complete network of tendons in both mutant and heterozygous control by E14.5. (B) Tendons of mutant pups appeared intact at birth, but by P3 lateral tendons disintegrated and were eventually eliminated (yellow arrowheads), whereas the majority of other tendons persisted with a substantial loss of the ScxGFP signal (white arrowheads). (C) Mutant pups appeared normal at birth but showed physical abnormalities including abducted paw and splayed limb (black arrowheads) by P3. (D-E) Substantial loss of ScxGFP signal was also detected in all tendons and related tissues. (D) Tail tendons and annulus fibrosus of the intervertebral disc (white arrowheads) in P7 pups. (E) Collateral ligaments of the metacarpophalangeal joint imaged in transverse section through the joints of heterozygous control and mutant pups at P7 (white arrowhead). Scale bar, 100 μm. Mutant: CKO, Heterozygous: Het.

Mutant tendon development was not perturbed through embryogenesis and mutant pups appeared normal at birth (Fig. 1C). However, by day 3 after birth (P3), mutant pups showed physical abnormalities that manifested in abducted paws, splayed limbs (Fig. 1C, black arrowhead) and severe movement limitations. Examination of forelimb tendons of P7 mutant pups using the tendon reporter ScxGFP revealed severe tendon disruptions. A few lateral tendons, e.g. the extensor carpi radialis longus tendon underwent fragmentation and disintegrated (Fig. 1B, yellow arrowhead and Fig. S1) whereas the majority of other tendons, notably the extensor digitorium communis tendons, retained structural integrity with a substantial loss of ScxGFP signal (Fig. 1B, white arrowhead). The substantial loss of ScxGFP signal was also detected in all tendons and related tissues, including hindlimb- and tail tendons, as well as ligaments and annulus fibrosus of the intervertebral disc (Fig. 1D,E). Movement limitations were exacerbated as mutant pups became older and all mutants died at or before P14. This phenotypic progression was observed in most mutant pups but intriguingly, in rare cases (∼2%) the mutant pups showed physical abnormalities and severe tendon phenotypes already at birth.

A closer examination of the mutant embryos identified the first indication of a tendon phenotype already at E16.5. The flexor carpi radialis tendons of mutant embryos were consistently torn by E16.5 (Fig. S2). Interestingly, this phenotype was highly reproducible while the patterning and development of other tendons in mutant embryos was not perturbed through embryogenesis. Moreover, expression of the prototypic tenocyte markers Scx, Tnmd and collagen I (Fig. 2A-D) and the development of the collagen matrix were not disrupted in any tendon of mutant embryos (Fig. 2E,F), including the flexor carpi radialis tendon before it snapped. A direct cause for the specific tear of the flexor carpi radialis tendon in mutant embryos was not identified to date.

**Fig 2.**
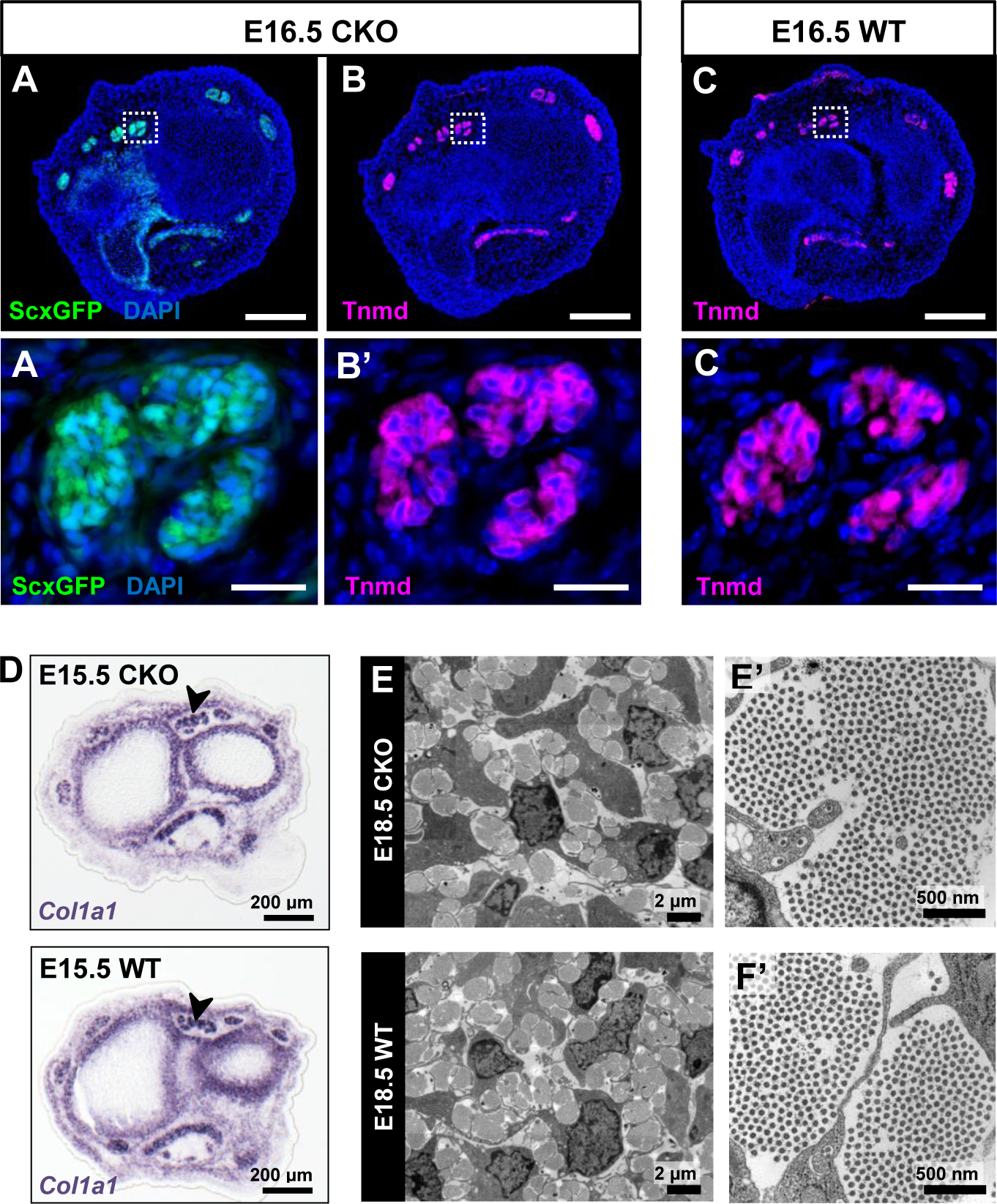
Tendon development in Tgfbr2;ScxCre mutant embryos was not perturbed through embryogenesis. (A) ScxGFP signal and (B) tenomodulin (Tnmd) immunofluorescence on transverse sections at wrist level of E16.5 mutant embryos demonstrate that the pattern and expression of prototypic tenocyte markers was not disrupted in mutant tendons. (C) Tnmd immunofluorescence in E16.5 wild-type tenocytes. (A’), (B’) and (C’) are higher magnifications of extensor digitorium communis tendons as boxed in (A), (B) and (C). (D) In situ hybridization for *Col1a1* on transverse sections of the forelimb from E15.5 mutant and wild-type littermates reveals that expression of the major matrix genes was not altered in mutant embryos (black arrowhead). (E,F) TEM images of tendons from forelimbs of E18.5 mutant and wild-type embryos reveals that organization and accumulation of the tendon extracellular matrix was not disrupted in the mutant. (E’,F’) Higher magnification views of (E) and (F) for direct visualization of the collagen fibers. Scale bars, 200 μm (A-C) and 20 μm (A’-C’). Mutant: CKO, Wild-type: WT.

Tendons are rich in collagen fibers that provide structural integrity to the tendons and transmit the forces generated by muscle contraction (Kannus, 2000). Since young mutant pups exhibited movement difficulties we first examined possible structural effects in the collagen matrix. The ultrastructure of mutant tendons that remained intact was therefore analyzed by transmission electron microscopy (TEM). Surprisingly, despite the functional defects starting around P3, collagen fibers in mutant tendons appeared organized and indistinguishable from those of wild-type (WT) littermates at this stage (Fig. 3). Apparent collagen degradation was observed only in older mutant pups (≥ P7) (Fig. 3), suggesting the disruption to the matrix of these tendons may be a secondary consequence of the cellular changes in these mutants and/or of their movement difficulties. Furthermore, epitenon, a monolayer of cells that engulf and define the boundary of the tendon (Kannus, 2000) (Fig. 3, black arrowhead), was gradually disrupted and in some cases was almost undetectable in older mutant pups (Fig. 3, white arrowhead), suggesting that loss of the tendon boundary is an additional feature of the phenotype in these mutants.

**Fig 3.**
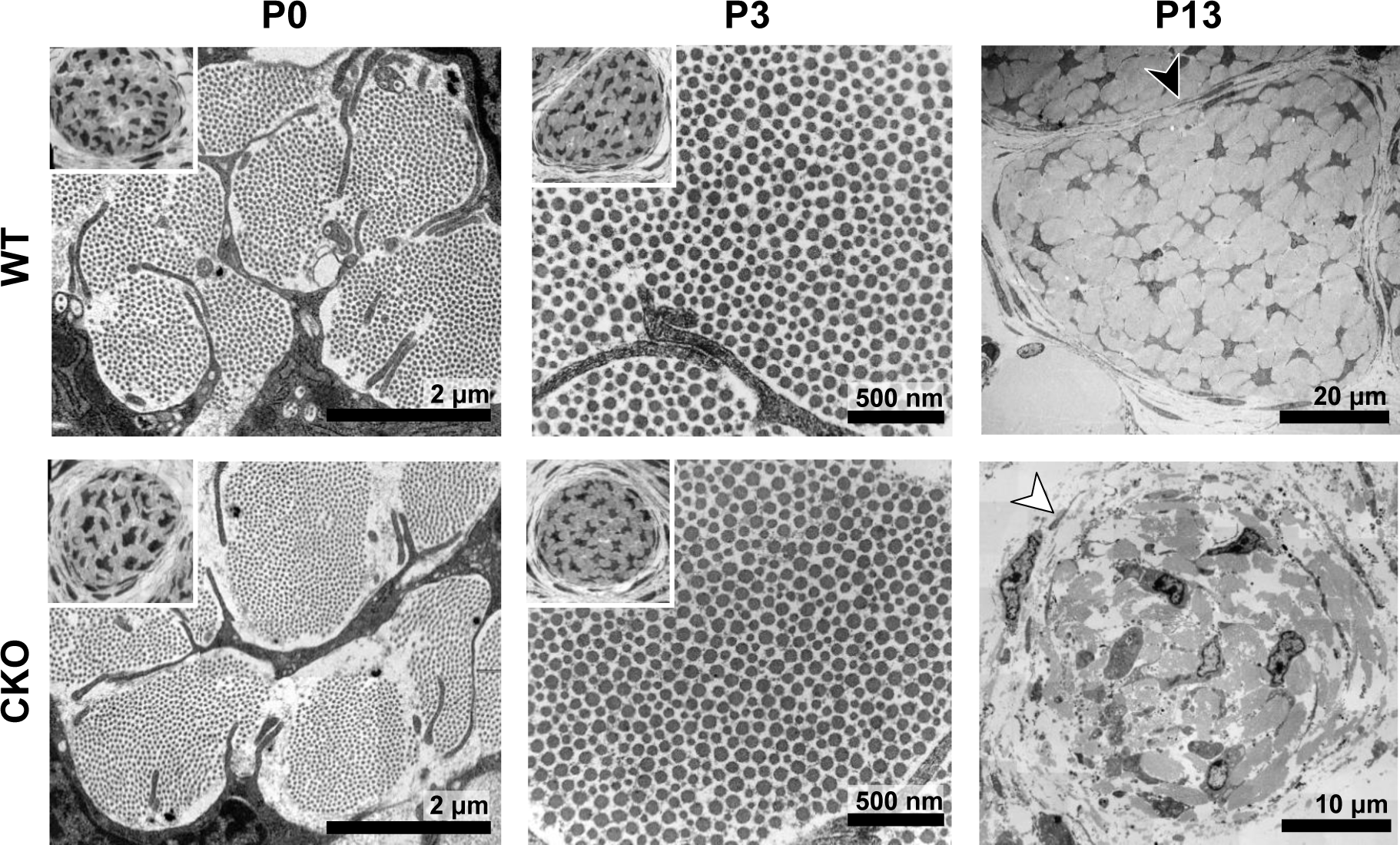
Tendon degeneration observed in Tgfbr2;ScxCre mutants at later postnatal stages. TEM images of tendons from forelimbs of mutant and wild-type littermates at P0, P3 and P13. Images were taken at high magnification for direct examination of structural details. Insets show transverse section TEM images of entire tendons at low-magnification (not to scale). Despite detectable functional defects starting around P3 in mutant pups, collagen matrix organization in mutant neonates was indistinguishable from that of their wild-type littermates. Apparent collagen degradation and disrupted epitenon structures (white arrowhead) were detected only in older mutant pups as shown here for P13 pups. Black arrowhead indicates epitenon in wild-type pups. Mutant: CKO, Wild-type: WT.

### Loss of the tendon cell fate in mutant tenocytes

As mentioned earlier, the ScxGFP signal in mutant tendons appeared patchy contrary to the smooth appearance of WT tendons (Fig. 1B), suggesting a disruption at the cellular level. To examine this phenotype at the cellular level we analyzed cross sections through the extensor communis tendons of P7 WT and mutant pups. In P7 WT pups, all tendon cells were positive for ScxGFP, *Tnmd* and *Col1a1* (Fig. 4A,C). Conversely, most cells in mutant tendons lost the ScxGFP signal and tendon marker gene expression (Fig. 4B, white arrowhead and 4C). Interestingly, some cells in mutant tendons retained ScxGFP signal and appeared rounded and enlarged from P3 onwards (Fig. 4B, yellow arrowhead). Further investigation suggests that these cells are newly recruited tendon cells. Analysis of this aspect of the mutant phenotype will be published in a separate manuscript (Tan et al., manuscript in preparation).

**Fig 4.**
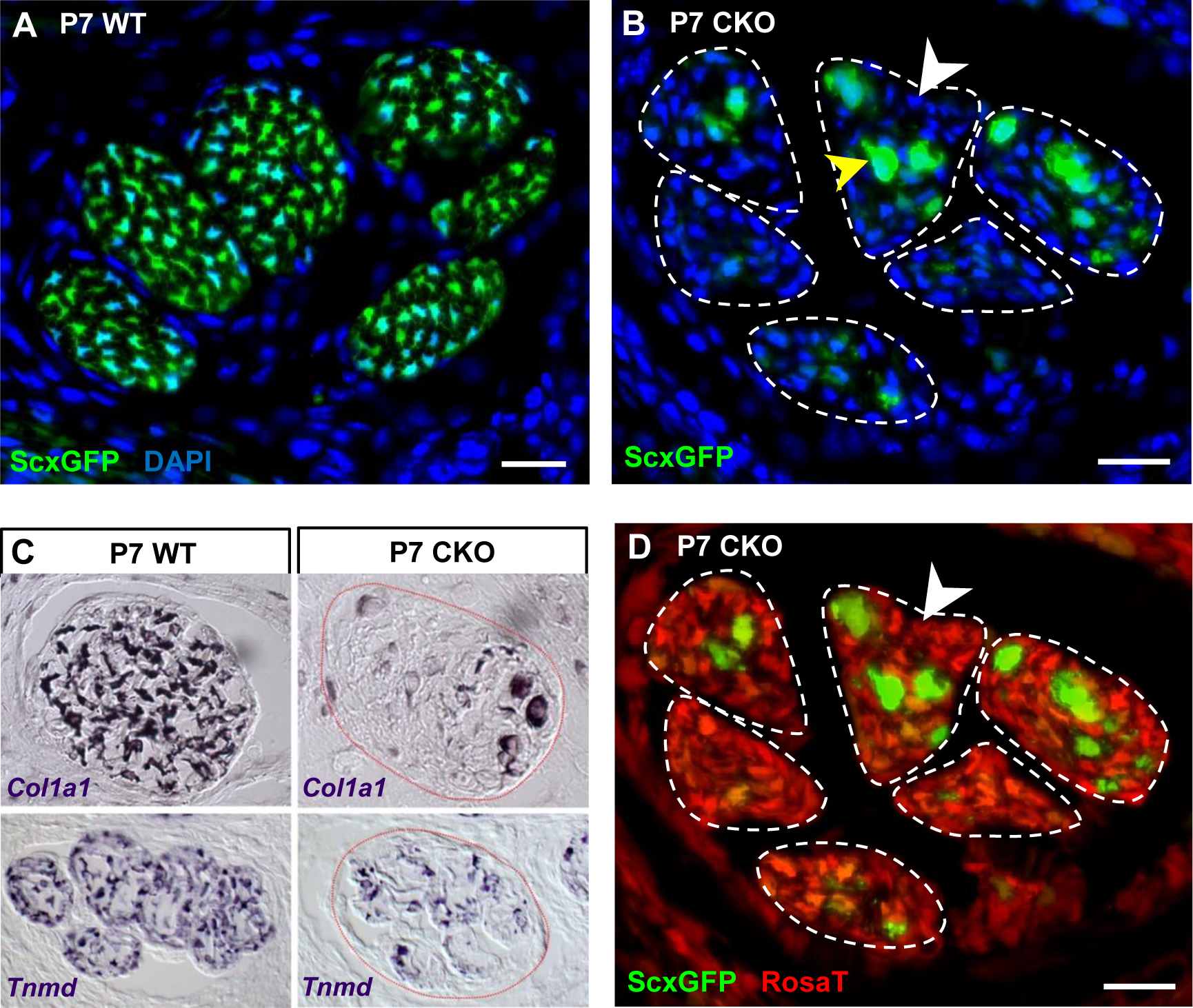
Deletion of Tgfbr2 in Scx-expressing cells (Tgfbr2;ScxCre) results in loss of tenocyte differentiation markers. (A-D) Transverse sections of extensor digitorium communis tendons of wild-type and mutant pups at wrist level. (A) In P7 wild-type pups, all tenocytes were positive for tendon reporter ScxGFP signal. (B) Conversely, most cells in P7 mutant tendons lost the ScxGFP signal (white arrowhead) whereas the cells positive for ScxGFP signal are newly recruited cells (yellow arrowhead; Tan et al, manuscript in preparation). (C) In situ hybridization shows that the mutant cells also lost gene expression of tendon markers *Col1a1* and *Tnmd* (images not to scale). (D) Lineage tracing using ScxCre shows that all ScxGFP-negative cells in (B) were positive for the Cre reporter Rosa26-tdTomato (RosaT; white arrowhead), proving that the ScxGFP-negative cells in mutant tendons were derived from the embryonic tenocytes. Dashed lines demarcate the mutant tendons. Scale bar, 20 μm. Mutant: CKO, Wild-type: WT.

The fact that most cells in the mutant tendons do not express tendon markers is surprising, since the cells in these tendons were functional tenocytes at embryonic stages as evidenced by tendon marker gene expression and by the development of a functional collagen matrix (Fig. 2). We next sought to determine if the ScxGFP-negative cells were indeed tendon cells that lost tendon gene expression or if the mutant tendons were simply repopulated by non-tenogenic cells. Using TUNEL assay we did not detect cell death in mutant tendons and the rate of tenocyte proliferation as examined by EdU assay was also not altered in these tendons (Fig. S3A,B), suggesting the cell population of mutant tendons was not altered. To directly determine if the cells in mutant tendons were tenocytes whose cell fate was altered, we took advantage of the fate mapping feature of the Rosa26-tdTomato (RosaT) Cre reporter system (Madisen et al., 2010). When the reporter is activated by ScxCre, expression of the RosaT reporter is restricted to the Scx-expressing cells and their progeny. We found that all ScxGFP-negative cells within mutant tendons were positive for the RosaT reporter (Fig. 4D, white arrowhead), proving that the cells in the mutant tendons were indeed derived from tenocytes. This result thus highlighted an unexpected reversibility for the tendon cell fate where it was possible for committed and functional tenocytes to lose their differentiation status.

To test if a continuous requirement for TGFβ signaling is essential for maintenance of the tendon cell fate, we next targeted Tgfbr2 either selectively in Scx-expressing cells or in all cells using the tamoxifen-inducible ScxCreERT2 (Howell et al., 2017) and RosaCreERT2 (Madisen et al., 2010) drivers respectively during different developmental stages (E13 and P0). Interestingly the cell fate of targeted cells was not disrupted in these mutants as evidenced by retention of differentiation marker expression (data not shown). This result suggests that tenocyte dedifferentiation in the Tgfbr2;ScxCre mutant tendons is dependent not simply on the loss of TGFβ signaling in these cells but also on additional features in these mutants associated with the specific spatial and temporal features of ScxCre activity.

### Mutant tenocytes acquired stem/progenitor features

Loss of cell differentiation marker can be the outcome of a few cellular processes, including most notably cell death, change of cell fate (transdifferentiation) or reversion to a less differentiated state (dedifferentiation) (Cai et al., 2007; Talchai et al., 2012; Tata et al., 2013). As aforementioned, we found no apparent cell death in mutant tendons (Fig. S3A). Using histological staining for the prototypic markers of osteocytes, adipocytes and chondrocytes we found that loss of tendon gene expression in the cells of mutant tendons was also not due to transdifferentiation (Fig. S3C), suggesting that the changes in mutant tendons may reflect a process of cellular dedifferentiation.

One hallmark of cellular dedifferentiation is the loss of differentiation markers, which is what we observed in mutant tendon cells. When cells dedifferentiate they also assume stemness features e.g. colony forming potential, and in most cases these cells also acquire expression of stem/progenitor cell markers (Sun et al., 2011; Tata et al., 2013; Nusse et al., 2018). To date, very little is known about the specific gene expression and cellular behavior of embryonic tendon progenitors. The only defined feature of these cells is the expression of the Scx tendon progenitor marker (Schweitzer et al., 2001), which was evidently lost in the mutant tendon cells. We therefore next directed our attention to similarities with stem/progenitor cells isolated from tendons (tendon-derived stem/progenitor cells) (Bi et al., 2007; Rui et al., 2010; Zhang and Wang, 2010; Mienaltowski et al., 2013) and with stem/progenitor cell markers reported in other studies (Blitz et al., 2013; Dyment et al., 2013; Tan et al., 2013; Runesson et al., 2015; Yin et al., 2016).

To test the colony-forming capacity of the mutant tendon cells, P7 mutant tendons were dissociated and FACS-sorted to collect ScxGFP-negative and RosaT-positive cells, which were then seeded at one cell per well in 96-well plates. As shown in Fig. 5A, about 1-2% of cells (ScxGFP-positive and RosaT-positive) isolated from tendons of P7 WT and heterozygous controls formed colonies in culture, similar to the frequency of colony forming cells reported in other studies (Bi et al., 2007; Rui et al., 2010). On the other hand, we found a significant 8-fold increase (*p*<0.01) in the frequency of colony forming cells in mutant tendons (Fig. 5A).

**Fig 5.**
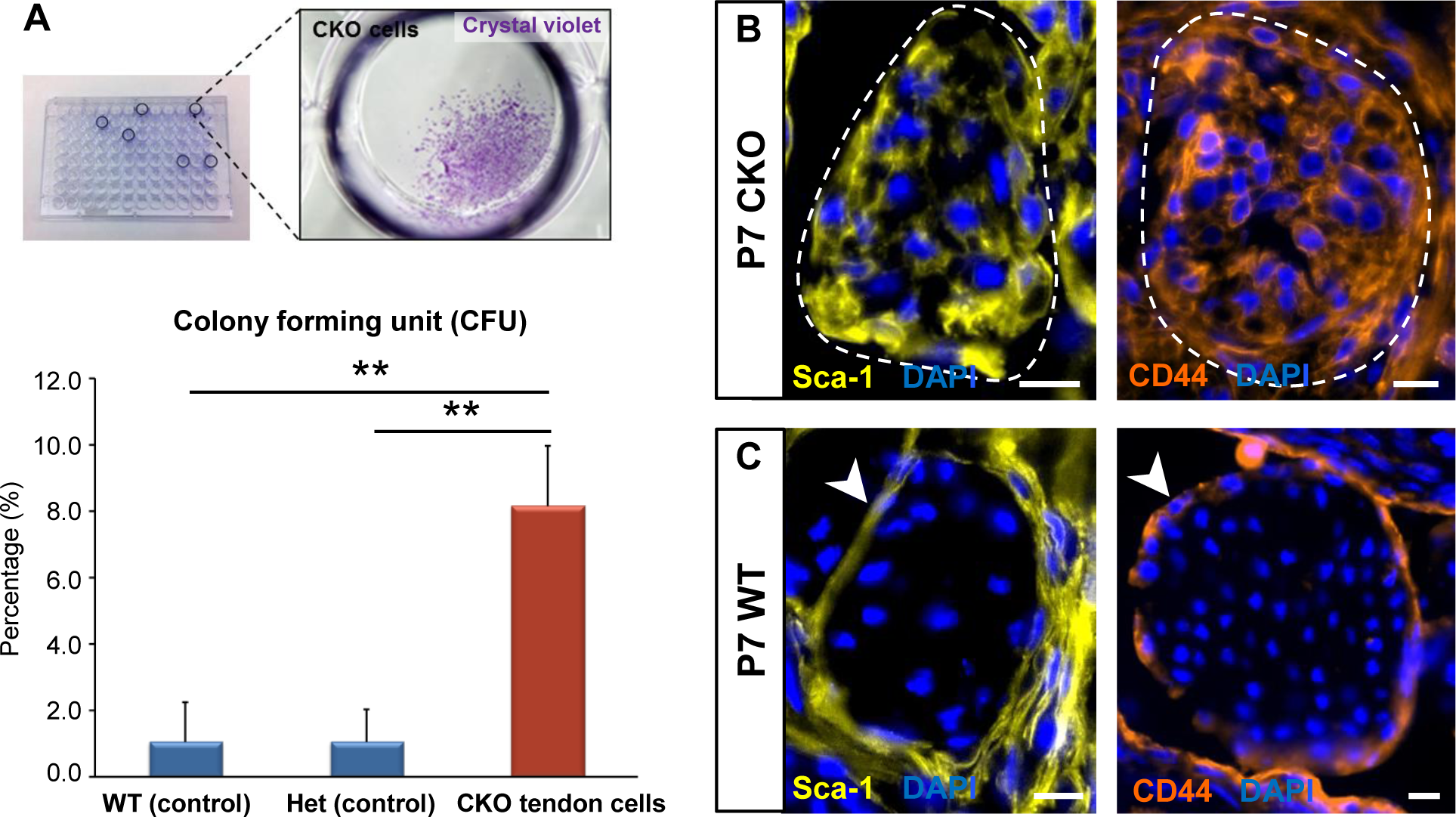
Tgfbr2;ScxCre mutant tenocytes acquired stem/progenitor features. (A) The colony-forming efficiency of P7 wild-type and heterozygous tenocytes as well as mutant tendon cells were evaluated by seeding one cell per well of the FACS-sorted cells in 96 well plates, and colonies formed were visualized with crystal violet staining. Mutant tenocytes exhibited significantly higher clonogenic capacity compared to wild-type and heterozygous controls. The results shown are mean ± SD (n=5-6, \*\**p*<0.01). (B) Immunofluorescence staining for stem/progenitor markers in transverse sections of mutant tendons shows that mutant tendon cells acquired in postnatal stages expression of stem cell antigen-1 (Sca-1) and CD44. (C) In wild-type littermate controls, expression of both markers was detected in epitenon (white arrowhead), but not in tenocytes. Dashed line demarcates the mutant tendon. Scale bars, 10 μm. Mutant: CKO, Wild-type: WT, Heterozygous: Het.

We next screened the mutant tendons for expression of stem/progenitor cell markers. We found that while Tgfbr2;ScxCre mutant tendon cells exhibited weak or negative expression of some progenitor markers e.g. CD90.2, Oct-3/4, Sox2, nucleostemin, alpha-SMA, nestin and Sox9 (data not shown), the mutant tendon cells gradually acquired expression of stem cell antigen-1 (Sca-1) and CD44 in postnatal stages (Fig. 5B). Notably, expression of Sca-1 was undetectable and CD44 was detected only in very few WT tendon cells, but surprisingly robust expression of these markers was detected in the epitenon (Fig. 5C, white arrowheads), a possible source of progenitor cells (Mendias et al., 2012; Dyment et al., 2013; Mienaltowski et al., 2013). The similarity of marker expression between the mutant tenocytes and epitenon cells therefore reinforces the notion that the mutant tenocytes acquired progenitor features.

Dedifferentiation is frequently associated with reversion to an earlier progenitor cell fate (Cai et al., 2007). We therefore next examined the expression of these markers during embryonic tendon development. At E12.5, when tendon progenitors are first detected (Pryce et al., 2009), expression of Sca-1 and CD44 could not be detected in ScxGFP-positive tendon progenitors (data not shown). At E14.5, at the onset of tendon cell differentiation, we found low or no expression of both markers in the differentiating tendon cells. Robust positive staining for both markers was however detected in the cells that surround the tendon at this stage, likely the precursors of the epitenon/paratenon (Fig. S4). Similar expression patterns were also found in mutant embryos (data not shown). These findings suggest that Sca-1 and CD44 are not markers for tendon progenitor *in vivo*, and possibly simply reflect a generic stemness state of the dedifferentiated mutant tendon cells.

Taken together, our findings show that mutant tendon cells acquired some stem/progenitor properties while losing their cell fate. It should be noted however that although these dedifferentiated tendon cells demonstrate some stem/progenitor properties, absence of TGFβ signaling in these cells might prevent them from acquiring the full spectrum of stemness or plasticity.

### Molecular profile of the dedifferentiated mutant tenocytes

We next performed single cell RNA sequencing analysis (scRNASeq) to establish a comprehensive profile of the cellular state and molecular changes in mutant tenocytes. A targeted retention of 2300-2600 cells from P7 WT- and mutant tendon was obtained, and the transcriptomes were analyzed using the 10X Genomics platform. Using unsupervised hierarchical clustering analysis, we identified two major clusters corresponding to WT tenocytes and mutant (dedifferentiated) cells in the respective samples. Expression of close to 1000 genes (mean UMI count ≥ 0.5, adjusted *p-*value ≤ 0.05) was identified in each of these clusters.

Pairwise comparison of the gene set between the P7 WT tenocyte and mutant cell clusters was next performed to determine changes in gene expression associated with tenocyte dedifferentiation. In total, expression of 186 genes was significantly different between the two cell populations (≥ 2-fold change and adjusted *p-*value ≤ 0.05), in which expression of 89 genes was upregulated and 97 genes was downregulated in the mutant tendon cells (Table S2). Almost 30% of the downregulated genes (29 genes) were identified in transcriptome analyses as tendon distinctive genes [(Havis et al., 2014) and our unpublished data]. Notably, the genes *Scx, Fmod, Tnmd, Pdgfrl, Col1a1, Col1a2, Col11a1 and Col11a2* were among the top 25 down-regulated genes in Tgfbr2;ScxCre mutant tendon cells (Table 1), further confirming the loss-of-cell fate phenotype in these cells. On the other hand, expression of the *Ly6a* gene (encoding Sca-1) was greatly enriched in P7 mutant cells, corroborating the IHC findings presented above (Table 2 and Fig. 5B). Moreover, we also found a significant increase in the expression of the *CD34* gene, another common marker for diverse progenitor cells. This observation was further confirmed at protein level, where positive immunostaining for CD34 was detected in mutant cells but not in normal tendon cells (Fig. 6A). Interestingly, the genes upregulated in the mutant cells included several genes (*Dpt*, *Anxa1*, *CD34*, *CD44*, *Mgp* and *Mfap5*) whose expression was previously reported to be enriched during embryonic tendon development (Havis et al., 2014). These findings thus do not only lend support to our notion that the mutant cells lost their differentiation state, but also suggest the possibility of induction of some developmental programs in these cells, a general feature in cellular dedifferentiation (Tata et al., 2013; Stocum, 2017; Nusse et al., 2018).

**Fig 6.**
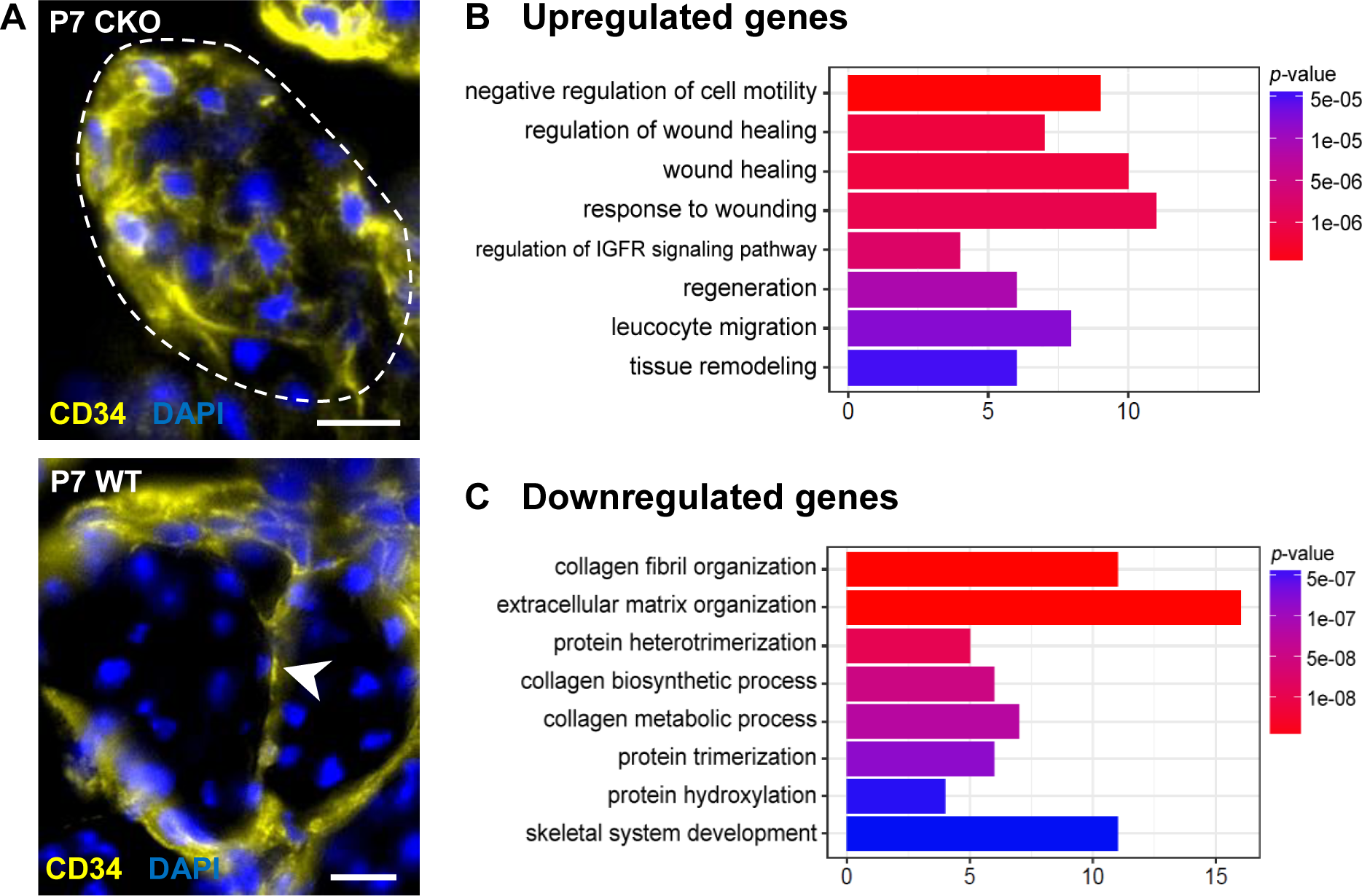
Molecular profile of the dedifferentiated mutant tenocytes. (A) Upregulated expression of *CD34* gene in P7 mutant tenocytes as revealed by scRNA-Seq analysis (see also Table 2) was determined using immunostaining. Transverse section of forelimb tendons shows that CD34 was indeed expressed by mutant tenocytes, while in wild-type controls CD34 was detected only in epitenon cells (white arrowhead). Dashed line demarcates the mutant tendon. (B,C) Gene ontology (GO) enrichment analysis in terms of biological processes associated with the (B) upregulated and (C) downregulated genes in P7 mutant compared with wild-type tenocytes. Selected GO terms are included in this figure, and genes annotated to the GO terms are available in Table S3. Scale bar, 10 μm. Mutant: CKO, Wild-type: WT.

**Table 1.**
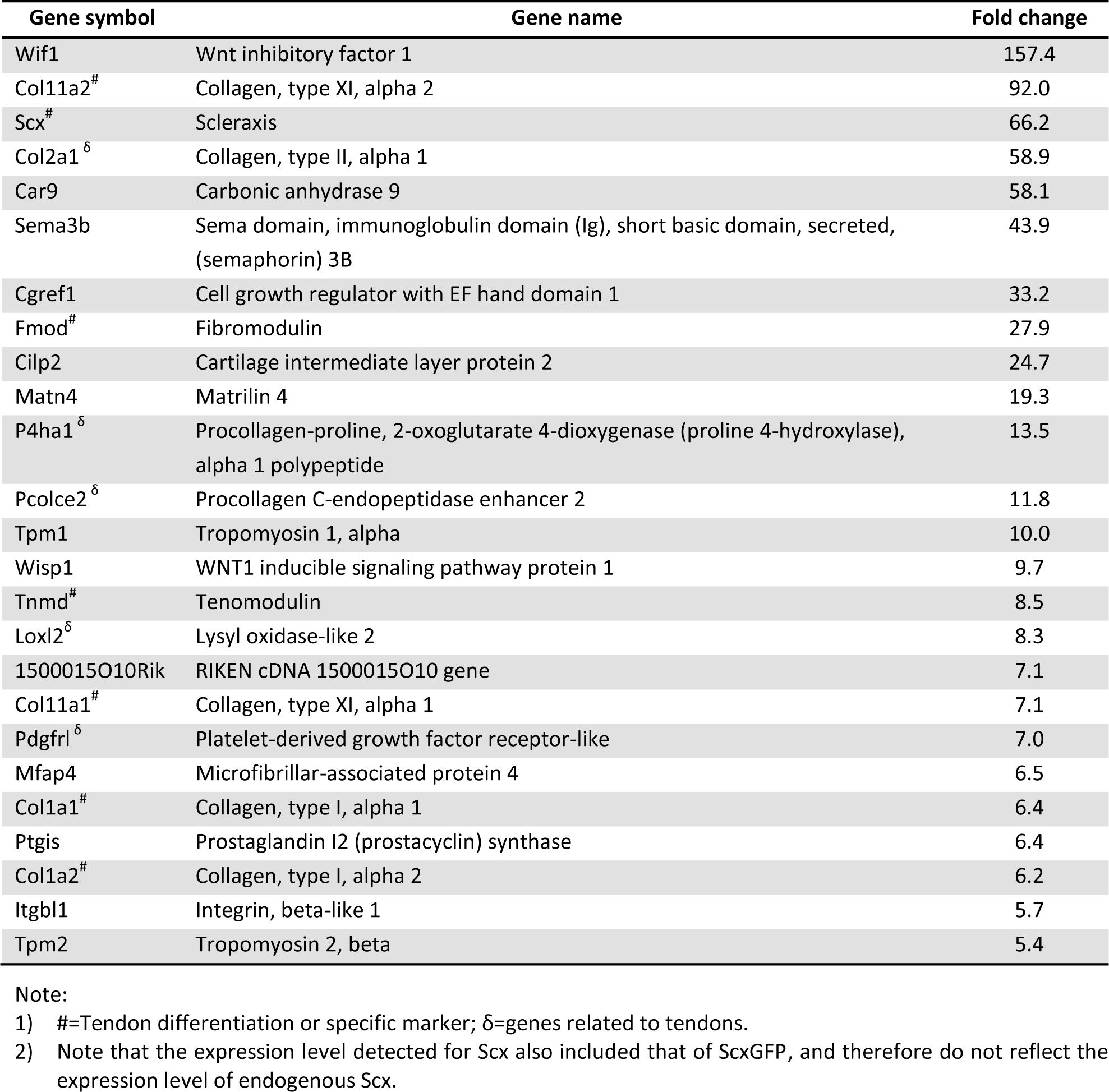
Top 25 downregulated genes (2-fold change, adjusted p<0.05) in P7 mutant cells compared with P7 wild-type tenocytes. See also Table S2 for a complete list of the downregulated genes.

**Table 2.** Top 25 upregulated genes (2-fold change, adjusted p<0.05) in P7 mutant cells compared with P7 wild-type tenocytes. See also Table S2 for a complete list of the downregulated genes.

| Gene symbol | Gene name | Fold change |
| --- | --- | --- |
| Dlk1 | Delta-like 1 homolog (Drosophila) | 137.9 |
| Serpine2 | Serine (or cysteine) peptidase inhibitor, clade E, member 2 | 118.2 |
| Dpt | Dermatopontin | 95.7 |
| Ly6a | Lymphocyte antigen 6 complex, locus A | 54.3 |
| H19 | H19 | 51.1 |
| Cd34 | CD34 antigen | 47.8 |
| Lum | Lumican | 36.6 |
| Lgmn | Legumain | 31.8 |
| Cxcl12 | Chemokine (C-X-C motif) ligand 12 | 26.1 |
| Mfap5 | Microfibrillar associated protein 5 | 22.5 |
| Ly6c1 | Lymphocyte antigen 6 complex, locus C1 | 21.7 |
| Igf2 | Insulin-like growth factor 2 | 21.4 |
| Serping1 | Serine (or cysteine) peptidase inhibitor, clade G, member 1 | 19.2 |
| Mgst1 | Microsomal glutathione S-transferase 1 | 18.3 |
| Aspn | Asporin | 15.9 |
| Mt1 | Metallothionein 1 | 15.4 |
| Mgst3 | Microsomal glutathione S-transferase 3 | 13.1 |
| Col3a1<sup>δ</sup> | Collagen, type III, alpha 1 | 13.0 |
| Postn | Periostin, osteoblast specific factor | 13.0 |
| Itm2a | Integral membrane protein 2A | 12.7 |
| Ptn | Pleiotrophin | 10.3 |
| Rps18-ps3 | Ribosomal protein S18, pseudogene 3 | 9.7 |
| Gsn | Gelsolin | 8.3 |
| Ifitm3 | Interferon induced transmembrane protein 3 | 8.2 |
| Col5a1<sup>δ</sup> | Collagen, type V, alpha 1 | 8.1 |

Note: $\delta$ =genes related to tendons.

To gain insights into biological functions activated in the P7 mutant cells, differentially expressed genes (DEGs) in these cells (Table S2) were further analyzed via GO enrichment tools clusterProfiler (Yu et al., 2012) and PANTHER Classification System (http://pantherdb.org/). Intriguingly, GO enrichment analysis revealed that one of the prominent biological changes observed in P7 mutant cells was upregulation of gene sets associated with wound healing (Fig. 6B and Table S3). These genes include protease inhibitors (*Serpine2, Serping1*), inflammatory mediator *Anxa1* and extracellular matrix (*Col3a1* and *Col5a1*). This finding suggests a possible role for tendon cells in the responses to pathological conditions, in line with findings reported by others (Dakin et al., 2015; Stolk et al., 2017; Schoenenberger et al., 2018). On the other hand, many biological processes downregulated in P7 mutant cells involved collagen synthesis and organization (Fig. 6C and Table S3). Since tendon biology is not annotated in most databases, changes in the collagen matrix, the most prominent structural component in tendons is the best indicator for the disruption of the tendon cell fate. Disruption of the collagen matrix in tendons was also detected in older mutant pups by ultrastructural analysis using TEM (Fig. 3).

Using PANTHER, we also investigated which protein classes were significantly altered in P7 mutant cells relative to WT tenocytes. Genes found to be most downregulated in mutant cells encode for receptors, signaling molecules, membrane traffic proteins and ECM (Table 3A). On the other hand, the upregulated genes in the mutant cells encode most prominently for proteins involved in nucleic acid binding, enzyme modulators, cytoskeletal protein, signaling molecules and transcription factors (Table 3B). Notably, expression of the activating protein 1 (AP-1) transcriptional complex, associated with numerous cellular processes including cell fate regulation (Hess et al., 2004), was significantly induced in mutant cells. Expression of both AP-1 components, i.e. the *Fos* and *Junb* genes was induced more than two fold, and the *Jun* gene was induced only slightly less than 2 fold. Moreover, the *Id3* gene encoding for a general bHLH transcription factor inhibitor was also induced. Due to its broad selection of targets, *Id3* was also implicated in numerous cellular processes including the regulation of cellular differentiation (Norton, 2000). A possible role for these transcriptional activities in tenocyte dedifferentiation will be addressed in future studies.

We next conducted PANTHER Pathway Analysis using different values of the filter parameter (mean UMI count and fold change) for enriching DEGs in P7 mutant cells. In general, we found that pathways that stood out as relevant for this study included integrin signaling, insulin/IGF, Wnt and inflammation mediated by chemokine and cytokine signaling pathways (Table 4). Insulin/IGF- and Wnt signaling are often implicated in cell proliferation and cell fate specification (Stewart and Rotwein, 1996; Sadagurski et al., 2006; Goessling et al., 2009; Salazar et al., 2016). It is interesting to note that their activation has also been associated with cellular dedifferentiation in skin, gut and neuron (Weber et al., 2003; Zhang et al., 2012; Perekatt et al., 2018). Further investigation is required to determine the specific roles of these signaling pathways in tenocyte dedifferentiation.

### Tenocyte dedifferentiation is dependent on cell autonomous loss of TGFβ signaling

Lastly we wanted to ask if tenocyte dedifferentiation in these mutants reflected a cell autonomous requirement for TGFβ signaling in tenocytes, or if it was the result of global changes that occurred in mutant tendons. To address this question we reactivated TGFβ signaling in isolated mutant tendon cells and determined the effects on tenocyte dedifferentiation. We previously found that transuterine injection of AAV viruses into embryonic limbs resulted in sporadic infection of limb tendons [unpublished data and (Huang et al., 2013)]. We therefore decided to address this question by injection of a Cre-activatable virus encoding an epitope tagged version of the receptor, AAV1-FLEX-Tgfbr2-V5 (Fig. 7A). Injection of this virus into embryonic mutant limbs would result in expression of Tgfbr2-V5 only in infected tendon cells due to the tendon-restricted activity of ScxCre in mutant embryos.

**Fig 7.**
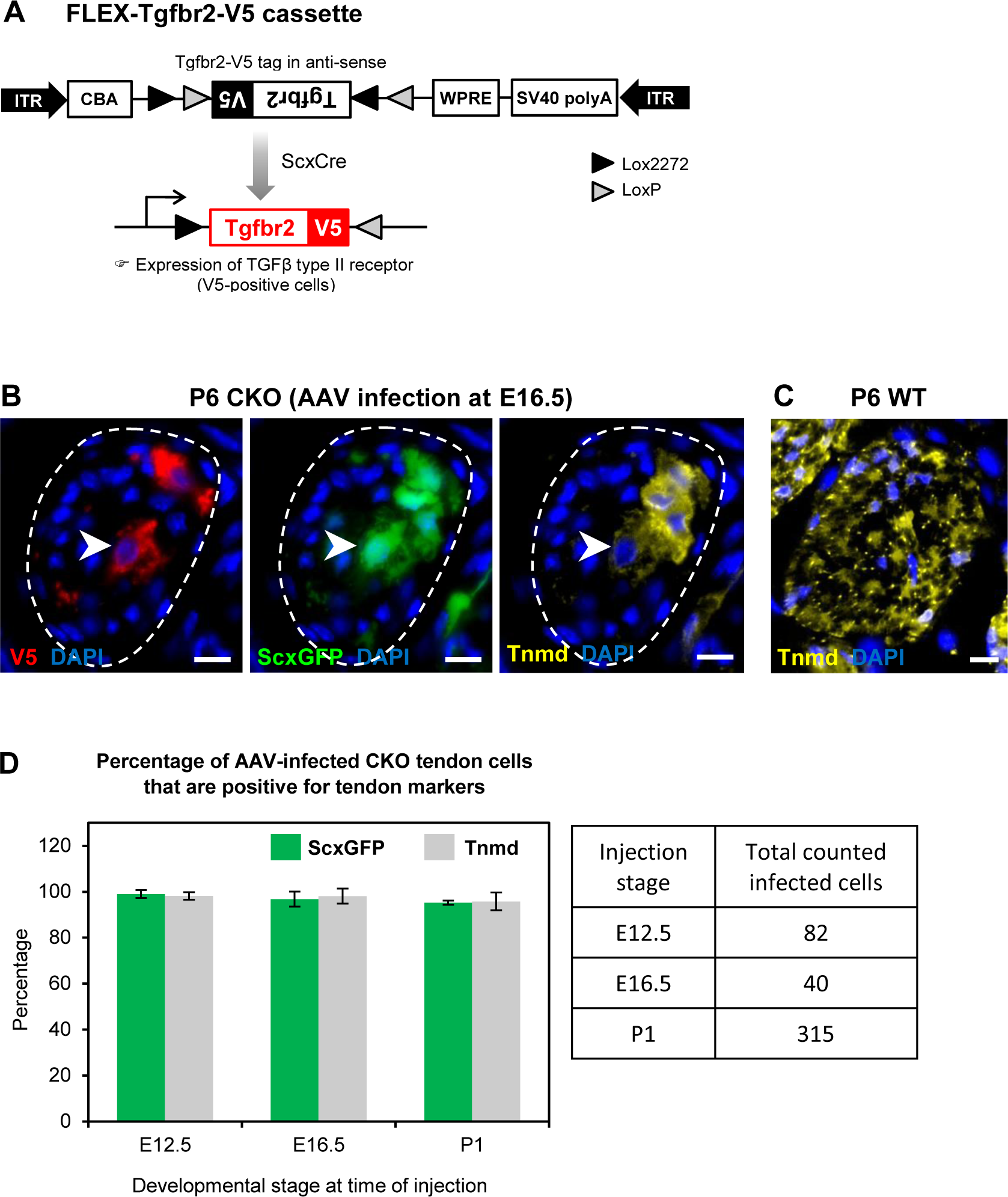
Tenocyte dedifferentiation is dependent on cell autonomous loss of TGFβ signaling. (A) AAV1-FLEX-Tgfbr2-V5 virus contains the reverse-complement sequence of Tgfbr2 with a C-terminal V5 epitope tag. Cre activity will lead to a permanent inversion of the cassette that will then express the V5-tagged TGFβ type II receptor. (B) Targeted expression of TGFβ type II receptor in E16.5 mutant tendon cells using the AAV1-FLEX-Tgfbr2-V5 prevented the loss of tendon markers in the infected tenocytes. The forelimb of E16.5 mutant embryos was infected with AAV1-FLEX-Tgfbr2-V5 virus and harvested at P6. Transverse forelimb sections were stained with antibodies for V5 (red) to detect AAV-infected cells and tenomodulin (Tnmd; yellow), a prototypic tendon marker expressed by (C) all tenocytes in the wild-type tendon at this stage. Dashed line demarcates the mutant tendon. (D) Quantification shows that about 95-98% of the AAV-infected (V5-positive) mutant tendon cells retained or re-expressed tendon differentiation markers after viral injection at different developmental stages (n = 3 pups for each stage). Scale bar, 10 μm. Mutant: CKO, Wild-type: WT.

AAV1-FLEX-Tgfbr2-V5 was injected into mutant limbs at two stages during embryonic tendon development: (a) E12.5 at the onset of ScxCre activity, ensuring that Tgfbr2-V5 expression will be activated in infected cells concurrent with the loss of the endogenous Tgfbr2, resulting in isolated Tgfbr2 expressing cells surrounded by mutant cells. (b) E16.5, before the onset of tenocyte dedifferentiation in mutant embryos. Interestingly, targeted re-expression of Tgfbr2-V5 in individual mutant tendon cell at both developmental stages was able to prevent the loss of tendon markers as observed in postnatal pups (Fig. 7B-D), suggesting a cell autonomous role for TGFβ signaling in maintenance of the tendon cell fate.

Recognizing that cell autonomous activity of Tgfbr2-V5 was sufficient to prevent dedifferentiation of mutant tenocytes, we next wanted to test if reactivation of TGFβ signaling in a dedifferentiating tenocyte could also reverse the process and rescue a tenocyte from dedifferentiation. Activity of ScxCre may be lost in the dedifferentiating tenocytes due to the loss of Scx expression and therefore of Scx enhancer driven expression of Cre in ScxCre mice. We therefore used in this case a virus encoding constitutive expression of Tgfbr2 in which the virus was tagged with a FLAG Tag (AAV1-Tgfbr2-FLAG). The virus was injected locally into P1 mutant limbs and the limbs were harvested at P7. We found that all infected mutant tendon cells expressed the tendon markers ScxGFP and tenomodulin (Fig. 7D and Fig. S5A), suggesting that reactivation of TGFβ signaling was indeed sufficient to rescue the dedifferentiated tenocytes. Taken together, these findings demonstrate that TGFβ signaling is sufficient to prevent and to rescue the loss-of-tendon cell fate in a cell-autonomous manner.

The constitutive expression of Tgfbr2-FLAG driven by the AAV1-Tgfbr2-FLAG virus ensured that the neonatal infection with this virus resulted in Tgfbr2-FLAG expression both within and outside of tendons. Notably, induction of tendon gene expression following activation of Tgfbr2-FLAG expression was detected only in dedifferentiated tenocytes and not in cells located outside of tendons (Fig. S5B). It was previously shown that TGFβ signaling is a potent inducer of ScxGFP and other tendon markers (Pryce et al., 2009; Maeda et al., 2011; Sakabe et al., 2018). This result reflects the fact that induction of tendon markers by TGFβ signaling is context-dependent and further indicates that the tenocytes in mutant pups have dedifferentiated to a state with tenogenic potential that retained the capacity to respond to TGFβ signaling.

Taken together these results highlight a surprising cell-autonomous role for TGFβ signaling in maintenance of the tendon cell fate. In Tgfbr2;ScxCre mutants tenocyte differentiation and function are normal during embryonic development but the tenocytes dedifferentiate in early postnatal stages. Tenocyte dedifferentiation is directly dependent on the loss of TGFβ signaling since retention or reactivation of the TGFβ receptor in isolated cells prevents or reverses the process of dedifferentiation. TGFβ signaling is thus essential for maintenance of the tendon cell fate.

## Discussion

In this study we find that the tendon cell fate requires continuous maintenance *in vivo* and identify an essential role for TGFβ signaling in maintenance of the tendon cell fate. To examine the different roles that TGFβ signaling may play in tendon development the Tgfbr2 gene was targeted in Scx-expressing cells (Tgfbr2;ScxCre mutant), ensuring disruption of TGFβ signaling in tendon cells. Mutant embryos appeared normal at birth and showed movement difficulties from early neonatal stages. Tendon formation and maturation was not affected in mutant embryos, but one flexor tendon snapped consistently at E16.5 and a few additional tendons disintegrated in early postnatal stages. Surprisingly, we find that in all other tendons the resident tenocytes lost tendon gene expression and dedifferentiated, assuming behavior and gene expression associated with stem/progenitor cells. While a direct loss of TGFβ signaling in individual tenocytes was not sufficient to cause tenocyte dedifferentiation, we found that tenocyte dedifferentiation could be reversed by reactivation of TGFβ signaling in mutant cells (Fig. 8). These results uncover an essential role for molecular pathways that maintain the differentiated cell fate in tenocytes and a key role for TGFβ signaling in these processes.

**Fig 8.**
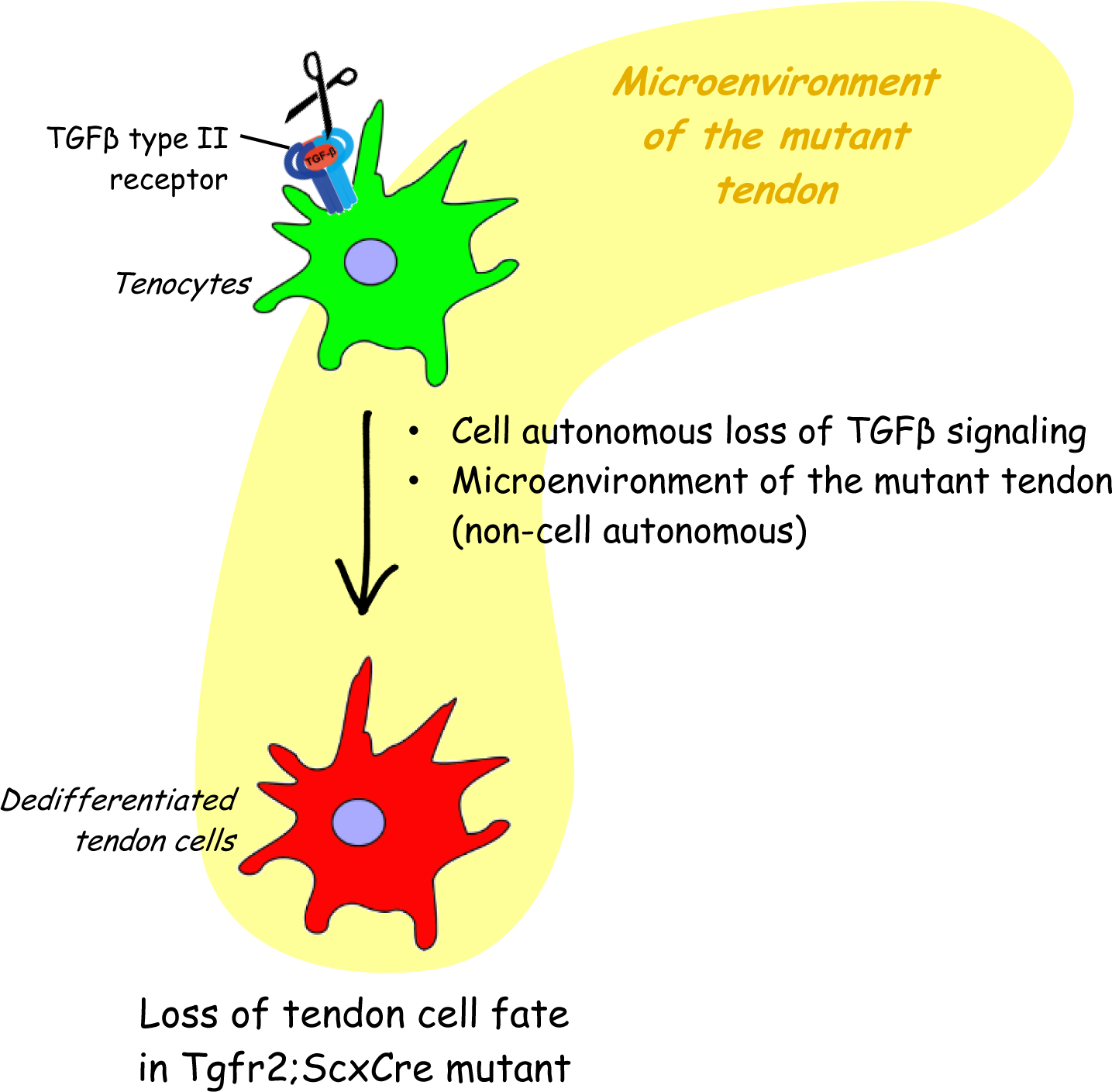
Proposed roles of TGFβ signaling in the maintenance of tendon cell fate. Targeted disruption of the TGFβ type II receptor by ScxCre resulted in tenocyte dedifferentiation in early postnatal stages. Tenocyte dedifferentiation was reversed by reactivation of TGFβ signaling in individual mutant cells, demonstrating a cell autonomous role for TGFβ signaling for maintenance of the cell fate. Conversely, direct loss of the TGFβ type II receptor using Cre drivers with different spatio-temporal features was not sufficient to cause tenocyte dedifferentiation, suggesting that mere loss of TGFβ signaling by tendon cells was not sufficient to cause dedifferentiation. We therefore propose that maintenance of the tendon cell fate is dependent on a combination of a cell autonomous function of TGFβ signaling and an additional, likely non cell-autonomous factor, e.g. the microenvironment of the tendon in the Tgbfbr2;ScxCre mutant.

Dedifferentiation has mostly been studied *in vitro* (Weinberg et al., 2007; Zhang et al., 2010; Pennock et al., 2015; Mueller et al., 2016; Guo et al., 2017; Nordmann et al., 2017) and there are only a handful of reported cases of dedifferentiation *in vivo* (Talchai et al., 2012; Tata et al., 2013; Zhang et al., 2019). It was therefore important to establish if the tenocytes of Tgfbr2;ScxCre mutants indeed dedifferentiated. Cellular dedifferentiation manifests in most cases by loss of features associated with the differentiated state and reversion to an earlier progenitor state within their cell lineage. In tendons of Tgfbr2;ScxCre mutants we indeed found that the tenocytes lost tendon gene expression and showed enhanced clonogenic potential. Moreover, the mutant tenocytes gained expression of the prototypic somatic stem/progenitor markers Sca-1, CD34 and CD44 (Holmes and Stanford, 2007; Sung et al., 2008; Hittinger et al., 2013; Sidney et al., 2014). Notably, of these stem/progenitor markers only Sca-1 and CD44 are also expressed at high levels in cultured tendon-derived stem/progenitor cells (Bi et al., 2007; Mienaltowski et al., 2013). Neither of these markers has so far been established as markers for tenocytes or for tendon progenitors. However, both expression of the CD34 and CD44 genes and expression of some additional signature genes identified in the dedifferentiated tenocytes by the scRNASeq analysis was previously shown to be significantly enriched in E14.5 mouse limb tendon cells when compared to cells from E11.5 (Havis et al., 2014). These observations suggest that some aspects of the embryonic tendon development program may be reactivated in dedifferentiated mutant tendon cells. Interestingly, we found that Sca-1, CD34 and CD44 are expressed in the wild-type epitenon/paratenon, thin layers of cells that surround the tendon and has been implicated as a possible source of stem/progenitor cells for tendons (Mienaltowski et al., 2013; Cadby et al., 2014). We further verified that mutant tendons are not repopulated by epitenon/paratenon cells since there is no evidence of elimination of the resident tenocytes by cell death.

Most studies of cellular dedifferentiation have focused on the regulation of this process *in vitro*. There is, however, evidence demonstrating this phenomenon *in vivo* especially in the context of pathological scenarios, as part of the regeneration process. One of the well-studied examples is limb regeneration in amphibians. Following limb amputation, cells near to the wound dedifferentiate to blastema, proliferate and eventually re-differentiate to replace all the components of the lost limb (McCusker et al., 2015). In zebrafish it has also been reported that following partial heart amputation, sarcomeres in mature cardiomyocytes disassembled, lost their differentiation gene expression profile and switched to embryonic hyperplastic growth to replace the missing tissues (Poss et al., 2002). Cellular dedifferentiation has also been observed in murine mature hepatocytes (Gournay et al., 2002), pancreatic β cells (Talchai et al., 2012) and skeletal muscle cells (Mu et al., 2011). More recently, Nusse and colleagues (Nusse et al., 2018) have shown that disruption of the mouse intestinal barrier, via either parasitic infection or cell death, led to reversion of crypt (epithelial) cells to a fetal-like stem cell state. Interestingly, expression of Sca-1 was highly induced in these cells, and when cultured the Sca-1 positive crypt cells exhibited characteristics of fetal intestinal epithelium including re-expression of fetal signature genes and loss of differentiated markers. The results presented in this study therefore suggest that a similar process may be activated in tenocytes as part of the regenerative process in response to pathology. Taken together, this growing body of evidence suggests that dedifferentiation may be a generalized cellular response to tissue damage that warrants further investigation. Moreover, these observations may also suggest that induction of Sca-1 may serve as a marker for a pathology-related dedifferentiation process. Intriguingly, Sca-1 positive cells were also found in the wound window in rat patellar tendon incisional injury model, but in this case it was not determined if Sca-1 expression was associated with dedifferentiation (Tan et al., 2013). Sca-1 expression has been identified on putative stem/progenitor cell populations in various tissues (Holmes and Stanford, 2007; Hittinger et al., 2013), but little is known about its biological function. It may therefore be interesting to examine whether Sca-1 functions as a stemness marker in dedifferentiated cells or if it also plays additional roles in cellular responses to pathological conditions.

Tenocyte dedifferentiation as observed in this study reveals an unexpected flexibility in the tendon cell fate where differentiated tenocytes can revert to a progenitor state under the mutant conditions Significantly, reintroduction of Tgfbr2 not only prevented tenocyte dedifferentiation when it was performed during embryogenesis but was also able to rescue the cell fate of dedifferentiated tenocytes when the virus was introduced after birth. This result suggests that TGFβ signaling may have a continuous role in protecting the differentiated tenocytes from dedifferentiation, identifying TGFβ signaling as a key regulator of tendon homeostasis. Moreover, these results also highlight the importance of the molecular pathways involved in maintenance of the differentiated cell fate not only for tissue homeostasis and function, but also for processes associated with tissue regeneration or with the onset and unfolding of pathology. Previous studies have implicated TGFβ signaling in cell fate maintenance in various tissues, e.g. preserving chondrocyte identity in cultures (Bauge et al., 2013) and suppressing intestinal cell dedifferentiation (Cammareri et al., 2017). While TGFβ signaling has been associated with different aspects of tendon biology (Pryce et al., 2009; Havis et al., 2016), to the best of our knowledge this is the first report of its role in maintenance of the tendon cell fate.

The fact that the mutant phenotype was caused by disruption of TGFβ signaling in tenocytes and the rescue of the tendon cell fate by virus mediated reintroduction of Tgfbr2 even to individual mutant cells provides direct evidence for a continuous and cell autonomous role for TGFβ signaling in maintenance of the tendon cell fate. However, targeting of Tgfbr2 using both tendon specific and ubiquitous inducible Cre drivers at different developmental stages did not result in tenocyte dedifferentiation, suggesting that tenocyte dedifferentiation in these mutants was dependent on specific spatial and temporal features of the ScxCre driver. These observations suggest that tenocyte dedifferentiation in these mutants may not merely be the result of the loss of intrinsic TGFβ signaling in tendon cells, but rather may be caused by an interplay between intrinsic loss of TGFβ signaling and additional external factors contributed by the environment of the mutant tendons, e.g. cell-cell or cell-matrix interaction.

The tendon phenotype of Tgfbr2;ScxCre mutants highlights a likely role for tenocyte dedifferentiation in regenerative processes in tendons and possibly also in the progression of tendon pathology. Uncovering the molecular pathways involved in this process may therefore be important for new strategies for treatments of tendon pathologies. The Tgfbr2;ScxCre mutants provide a unique opportunity to analyze these pathways, and the experimental approaches employed in this study may be developed into an experimental paradigm for molecular dissection of this process. Briefly, transcriptional and epigenetic analyses of the mutant tenocytes through the dedifferentiation process can provide a landscape of the molecular changes that initiate and drive the dedifferentiation process. Promising candidates can then be tested using the AAV-mediated cell fate rescue experiments to identify genes or groups of genes that can protect the tenocytes from dedifferentiation to establish the molecular process of cellular dedifferentiation. Of particular interest will be the early molecular changes in the mutant tenocytes that drive and promote the onset and progression of tenocyte dedifferentiation.

Our findings underscore the fact that the tendon cell fate requires continuous maintenance and that it is not an irreversible state, a long-standing biological dogma that has been challenged by recent research (Takahashi and Yamanaka, 2006; Ladewig et al., 2013). Nevertheless, it is important to recognize that the dramatic cell fate changes in Tgfbr2;ScxCre mutant happens in the context of a genetic modification. The occurrence of such phenomenon *in vivo* might not be a simple direct outcome of changes to TGFβ signaling. Most importantly, while the initiating events for tenocyte dedifferentiation may vary in different scenarios, it is likely that the molecular events that drive the dedifferentiation process downstream of the initiation event are similar or related. Uncovering these pathways in this experimental system may therefore facilitate the analysis of such processes in various other contexts.

## Materials and Methods

### Mice

All mouse work was performed in accordance to the guidelines issued by the Animal Care and Use Committee at Oregon Health & Science University (OHSU). Floxed TGFβ type II receptor (Tgfbr2^f/f^) mice (Chytil et al., 2002) were crossed with mice carrying the tendon deletor Scleraxis-Cre recombinase (ScxCre) (Blitz et al., 2013), to disrupt TGFβ signalling in tenocytes. All mice in this study also carried a transgenic tendon reporter ScxGFP (Pryce et al., 2007), and a Cre reporter Ai14 Rosa26-tdTomato (RosaT) (Madisen et al., 2010) for the lineage tracing of Scx-expressing cells. For embryo harvest, timed mating was set up in the afternoon, and identification of a mucosal plug on the next morning was considered 0.5 days of gestation (E0.5). Embryonic day 14.5 to postnatal day 13 (E14.5-P13) limb tendons were used for analysis. Mouse genotype was determined by PCR analysis of DNA extracted from tail snip using a lysis reagent (Viagen Biotech, Cat 102-T) and proteinase K digestion (55°C, overnight).

### Transmission electron microscopy (TEM)

Skinned mouse forelimbs were fixed intact for several days in 1.5% glutaraldehyde/1.5% formaldehyde, rinsed, then decalcified in 0.2M EDTA with 50mM TRIS in a microwave (Ted Pella, Inc.) operated at 97.5 watts for fifteen 99 min cycles. Samples were fixed again in 1.5% glutaraldehyde/1.5% formaldehyde with 0.05% tannic acid overnight, then rinsed and post-fixed overnight in 1% OsO_4_. Samples were dehydrated and extensively infiltrated in Spurr’s epoxy and polymerized at 70°C (Keene and Tufa, 2018). Ultrathin sections of tendons of interest were cut at 80 nm, contrasted with uranyl acetate and lead citrate, and imaged using a FEI G20 TEM operated at 120 kV with montages collected using a AMT XR-41 2 x 2K camera. The acquired images were stitched using ImageJ software (https://imagej.nih.gov/ij/) (Preibisch et al., 2009). Three pups per time point were harvested for TEM analysis.

### In situ hybridization and histological staining

Dissected forelimbs were fixed with 4% paraformaldehyde in PBS, decalcified in 5 mM EDTA (1-2 weeks at 4°C) and incubated with a 5-30% sucrose/PBS gradient. The tissues were then embedded in OCT (Tissue-Tek, Inc), sectioned at 10-12 µm using a Microm HM550 cryostat (Thermo Scientific, Waltham, MA) and mounted on Superfrost plus slides (Fisher). In situ hybridization was performed as previously described (Murchison et al., 2007).

For immunofluorescence staining, sections were air-dried, rinsed thrice with PBS and blocked with 2% BSA and 2% normal goat serum in PBS for 1h at RT. The sections were then incubated overnight at 4°C with specific primary antibody as listed in Table S1. This was followed by incubation with the matching Cy3- or Cy5-conjugated secondary antibody (Jackson ImmunoResearch; diluted at 1:250 or 1:400) in PBS containing 2% normal goat serum for 1h at RT. DAPI (4′,6-diamidino-2-phenylindole, dihydrochloride; Life Technologies) was used to counterstain cell nuclei. Immunolabelled sections were mounted in Fluorogel (Electron Microscopy Sciences, PA; Cat 17985-10) and visualized using a Zeiss ApoTome microscope. A washing step with PBS containing 0.1% Triton-X 100 was performed after the change of antibodies. Controls included omission of primary antibodies.

For examination of cell death and proliferation, TUNEL and EdU assays were performed using Click-iT EdU (Life Technologies) and In Situ Cell Death Detection (Roche) kits, respectively, following manufacturer’s instructions. For all studies, sections from 2-4 pups were examined to ensure reproducibility of results.

### Isolation and culture of tendon-derived stem/progenitor cells

Mice at P7 were used for tendon progenitor cell isolation using a protocol modified from that in Mienaltowski et al (Mienaltowski et al., 2013). Briefly, both forelimbs and hindlimbs were harvested from euthanized mice, skinned and exposed to 0.5% collagenase type I (Gibco, Cat 17100-017) and 0.25% trypsin (Gibco, Cat 27250-018) in PBS for 15 min at 37°C with gentle shaking. The surfaces of tendons were then scraped carefully with a pair of forceps to remove epitenon/paratenon cells. The middle portion of tendons was then harvested, cut into small pieces and tendon cells were released by digestion for 30 min at 37°C with gentle shaking in a solution of 0.3% collagenase type I, 0.8% collagenase type II (Cat 17101-015), 0.5% trypsin and 0.4% dispase II (Cat 17105-041) (all from Gibco). The released cells were strained with a 70-μm cell strainer (BD Falcon, Cat 352350) and collected by centrifugation for 5 min at 300 g. The cells were then resuspended in PBS with 1% BSA, and fluorescence-activated cell sorting (FACS) was used to separate the cells for colony-forming assay.

### Colony-forming unit (CFU) assay

CFU assay was used to examine the self-renewal potential of cells (Bi et al., 2007). The enzymatically-released WT and heterozygous tenocytes as well as dedifferentiated mutant tendon cells (i.e. ScxGFP-negative and RosaT-positive cells) were sorted by FACS and plated at one cell per well in a 96-well plate using a BD Influx^TM^ cell sorter (BD Bioscience, USA). About 10-12 days into the culture, the colonies were fixed in 4% paraformaldehyde (10 min, RT), stained with 0.5% crystal violet for 30 min, and rinsed twice with water. Percentage of colony-forming unit was calculated as: Number of wells with colonies ÷ 96 x 100. Each data point represents the mean of duplicate plates from 3-5 separate experiments. Each experiment represents limb tendons collected from 2-4 pups.

### Re-expression of Tgfbr2 in mutant cells using adeno-associated virus (AAV) vector

FLAG or V5 epitope tag sequences were added at the C-terminus of the murine TGFbR2 Consensus Coding Sequence (CCDS23601). The Tgfbr2-FLAG (*Tgfbr2-FLAG*) and reverse-compliment Tgfbr2-V5 (*FLEX-Tgfbr2-V5*) insert sequences were synthesized and subcloned by GenScript into an AAV vector. The FLEX backbone vector (Atasoy et al., 2008) was purchased from AddGene and modified. Vectors were then packaged into AAV1 capsid, purified, and titered by the OHSU Molecular Virology Support Core. AAV insert expression was under the control of a chicken beta-actin (CBA) promoter and an SV40 polyadenylation sequence. All experimental procedures were evaluated and approved by the institutional Animal Care and Ethics Committee.

Re-expression of Tgfbr2 in embryos was done by delivery of AAV-FLEX-Tgfbr2-V5, a Cre-dependent expression cassette, specifically to Tgfbr2^f/-^;ScxCre mutant tendon cells. Transuterine microinjection of the viral vector into embryos was performed according to a published protocol (Jiang et al., 2013). Briefly, a laparotomy was performed on anesthetized pregnant females to expose the uterus. The left wrist field of the forelimb bud of each embryo was injected with ∼2 µl of concentrated viral inoculum (3.8 x 10^13^ vg/ml) using a borosilicate glass capillary pipette (25-30 µm outer diameter and 20 degree bevel). The abdominal and skin incisions were closed with resorbable sutures. The dams were recovered overnight with supplementary heating and then returned to main colony housing.

For postnatal re-expression of Tgfbr2, ∼10 µl of AAV-Tgfbr2-FLAG inoculum (4.1 x 10^12^ vg/ml) was injected subcutaneously into the left forelimb of P1 pups using an 8 mm x 31G BD Ultra-Fine^TM^ insulin syringe and needle (Becton Dickinson and Company, NJ, USA). For both experiments, forelimbs from P5 to P7 mutant pups (n = 3 pups for each stage) were harvested, fixed, cryosectioned and examined for expression of tendon differentiation markers in infected tendon cells.

### Single-cell RNA sequencing (scRNA-seq) and data analysis

Tendons were collected and pooled from both forelimbs and hindlimbs as described above from two pups with the omission of tissue-scraping step. The enzymatically-released cells were centrifuged, resuspended in α-MEM with 5% FBS and submitted to the OHSU Massively Parallel Sequencing Shared Resource (MPSSR) Core facility. scRNA-seq analysis was then performed using the 10x Genomics Chromium™ Single Cell 3’ Reagent Kits and run on a Chromium™ Controller followed by sequencing using the Illumina NextSeq^®^ 500 Sequencing System (Mid Output), as per the manufacturer’s instructions (10x Genomics Inc, CA; Illumina Inc, CA).

Sequencing data processing and downstream analysis were performed using Cell Ranger version 2.0 (10x Genomics, CA) (Zheng et al., 2017) with the default settings. Briefly, sequencing reads were aligned to the mm10 genome and demultiplexed and filtered using total UMI count per cell to generate the gene barcode matrix. Principle component analysis was performed and the first ten principle components were used for the tSNE dimensional reduction and clustering analysis. Cells were clustered using K-means clustering. For each cluster, genes with an average UMI count ≥ 0.5, fold change ≥ 1.5 and *p*-value ≤ 0.05 were identified as signature genes for each cluster. Gene Ontology (GO) enrichment analysis (clusterProfiler) (Yu et al., 2012) and the PANTHER Classification System (http://pantherdb.org/) were used to elucidate the biological process and signaling pathway associated with individual gene. Enriched canonical pathways were defined as significant if adjusted *p*-values were < 0.05.

### Statistical analysis

Unless stated otherwise, all graphs are presented as mean ± standard deviation (SD). Student’s t-tests were performed to determine the statistical significance of differences between groups (n ≥ 3). A value of *p*<0.05 is regarded as statistically significant.

## Acknowledgements

The authors thank Dr Elazar Zelzer (Department of Molecular Genetics, Weizmann Institute of Science, Israel) for critical reading of the manuscript. We are grateful to staff from MPSSR and FACS core facilities, OHSU particularly Dr Robert Searles, Mrs Amy Carlos and Dr Miranda Gilchrist for their excellent technical assistance. This work was funded by NIH (R01AR055973, RS; R01DC014160, JVB) and Shriners Hospitals for Children (SHC 5410-POR-14). G.K.T was supported by Research Fellowship from Shriners Hospital for Children.

## Supplementary files

**Fig S1.**
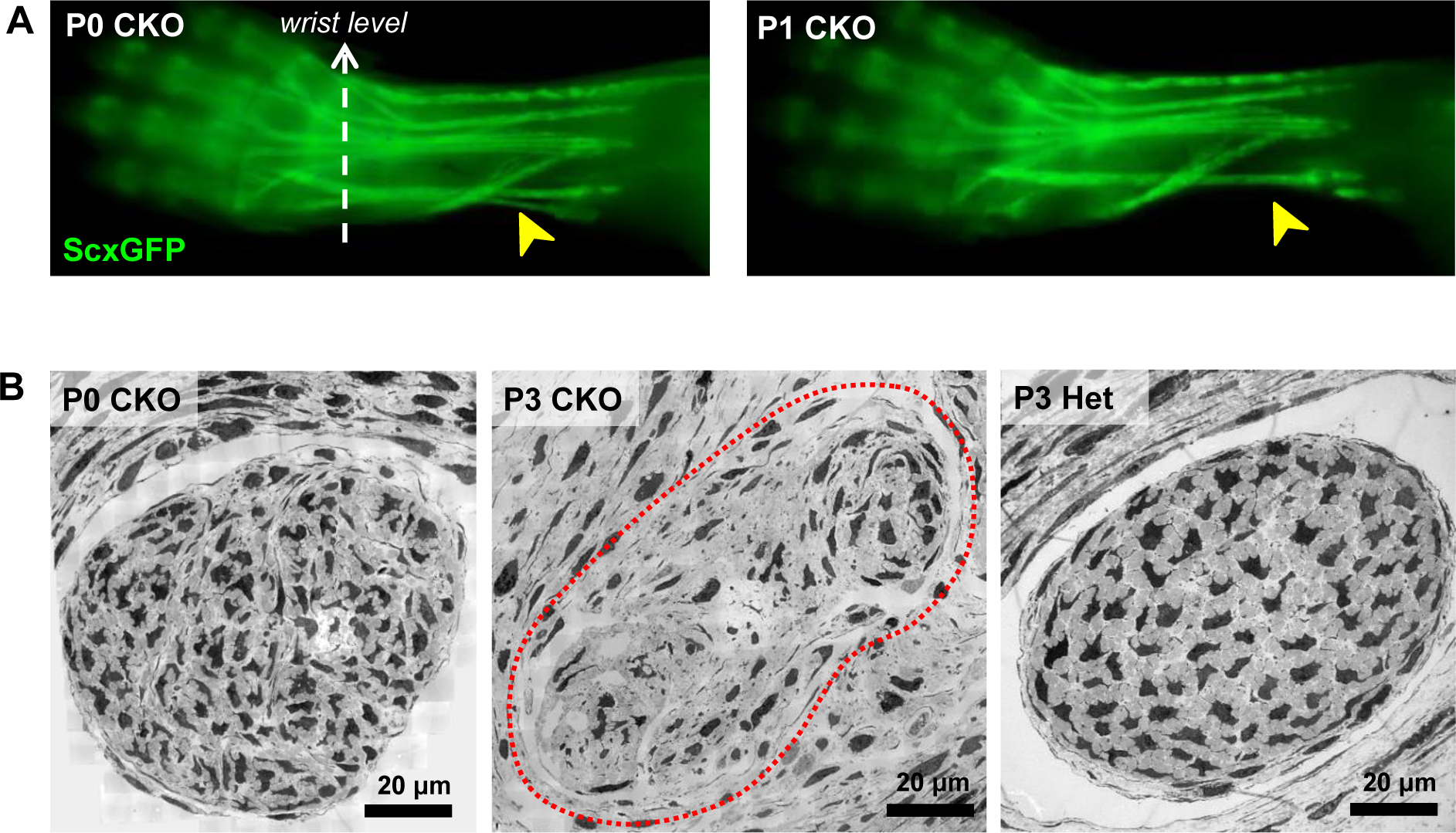
Fragmentation and elimination of lateral tendons in mutant neonates. (A) Rapid disruption of lateral extensor tendons in neonatal stages of mutant pups revealed by examination of forelimb tendons using the tendon reporter ScxGFP. The extensor carpi radialis longus tendon (yellow arrowheads) is present in a mutant pup at P0 but lost in a P1 mutant. (B) TEM images of the extensor carpi radialis longus tendon at wrist level. The mutant tendon shows signs of fragmentation already at P0, and by P3 the tendon appears disintegrated accompanied by complete loss of the epitenon and structural definition of the tendon circumference. The red dotted line in (B) demarcates the mutant tendon. Mutant: CKO, Heterozygous: Het.

**Fig S2.**
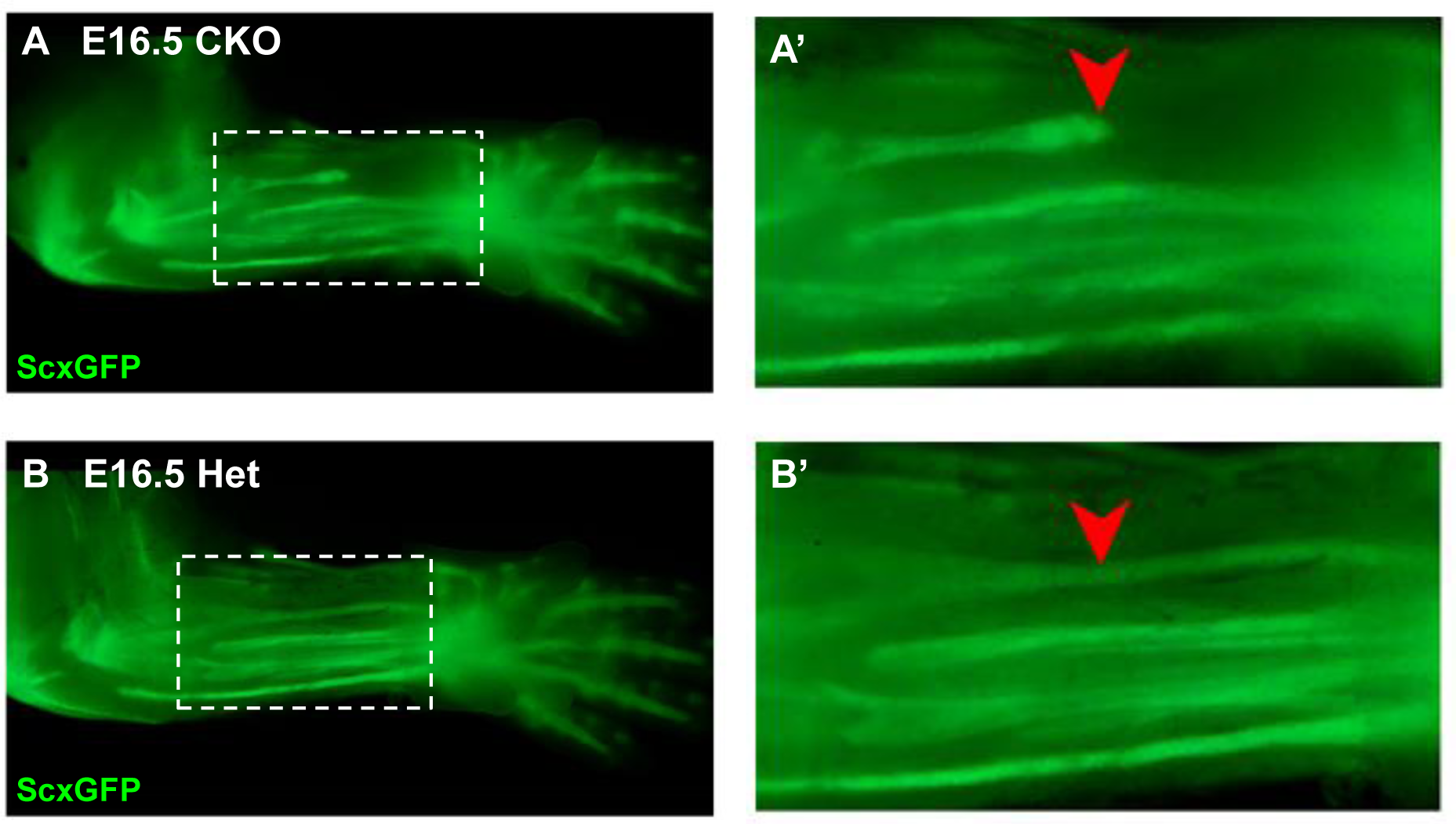
Disruption of the flexor carpi radialis tendon in mutant embryos. Examination of flexor tendons in E16.5 (A) mutant and (B) heterozygous littermates using the tendon reporter ScxGFP. Boxed regions in (A) and (B) are shown enlarged in (A’) and (B’). While most tendons appeared normal in mutant embryos, starting at E16.5 the flexor carpi radialis tendon (red arrowheads) was consistently torn in mutant embryo. Mutant: CKO, Heterozygous: Het. Figures not to scale.

**Fig S3.**
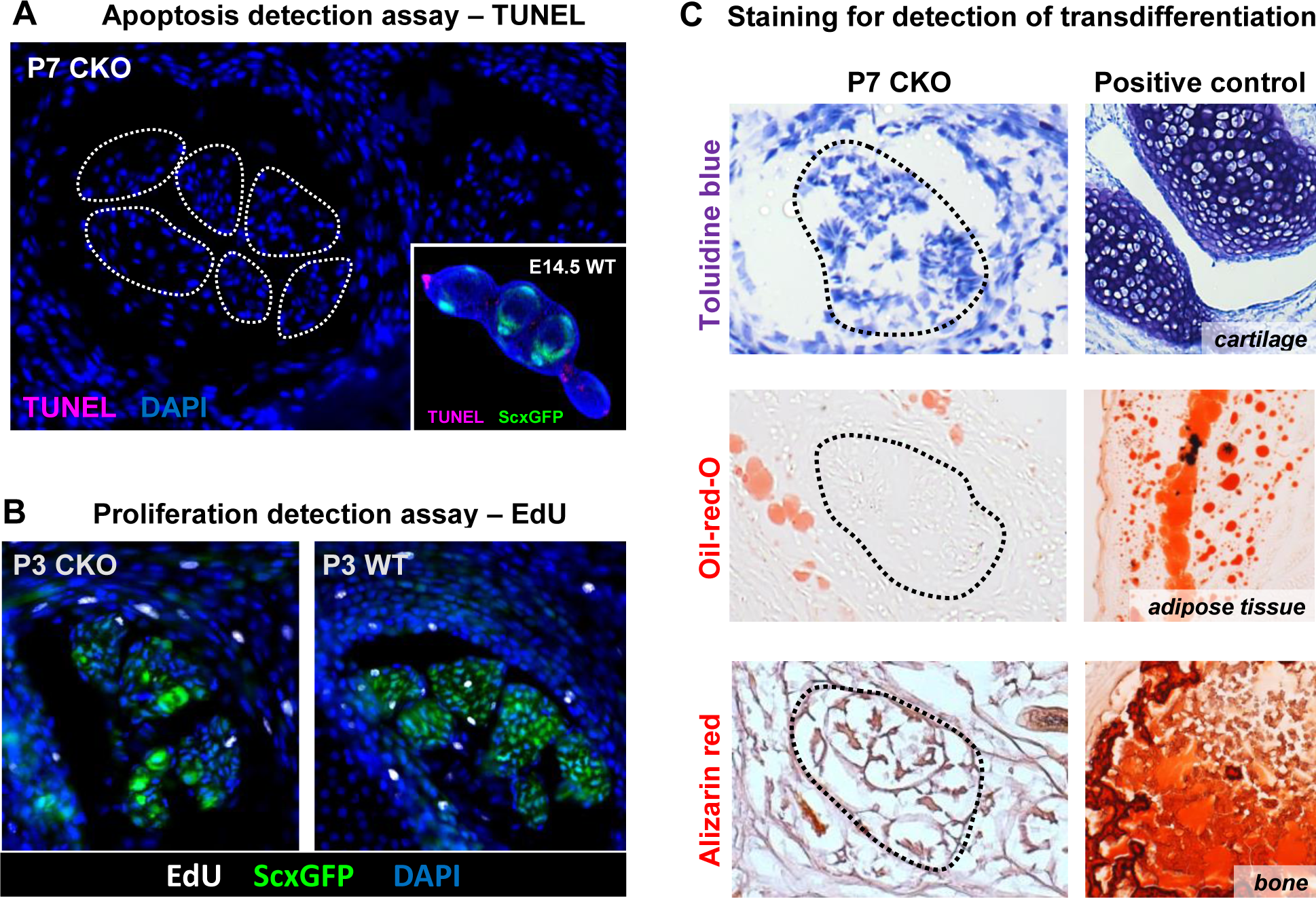
Evaluating cell death, proliferation and transdifferentiation in mutant tendons. (A) TUNEL assay did not detect significant cell death in mutant tendons. The extensor digitorium communis tendons are demarcated in a transverse section of P7 mutant forelimb. Inset in (A) shows a transverse section of E14.5 forelimb paw that serves as a positive control for TUNEL staining. As expected, cell death is detected only at the distal edge of the autopod, but not in tendons (ScxGFP) at this stage. (B) EdU labeling of proliferating cells in transverse sections of the forelimb from P3 pups. The rate of proliferation was also not altered in mutant tendons compared with the wild-type littermates. The pups were injected i.p. with 100 μg of EdU in PBS and tissues were harvested two hour post injection. (C) Histological staining for the prototypic markers of chondrocytes (toluidine blue), osteocytes (alizarin red) and adipocytes (oil-red-o) showed that the loss of tendon markers in mutant tenocytes was also not due to transdifferentiation. The positive control tissues for the respective staining are cartilage, adipose tissue and bone from the same section. Dotted lines demarcate tendons. Mutant: CKO, Wild-type: WT. Figures not to scale.

**Fig S4.**
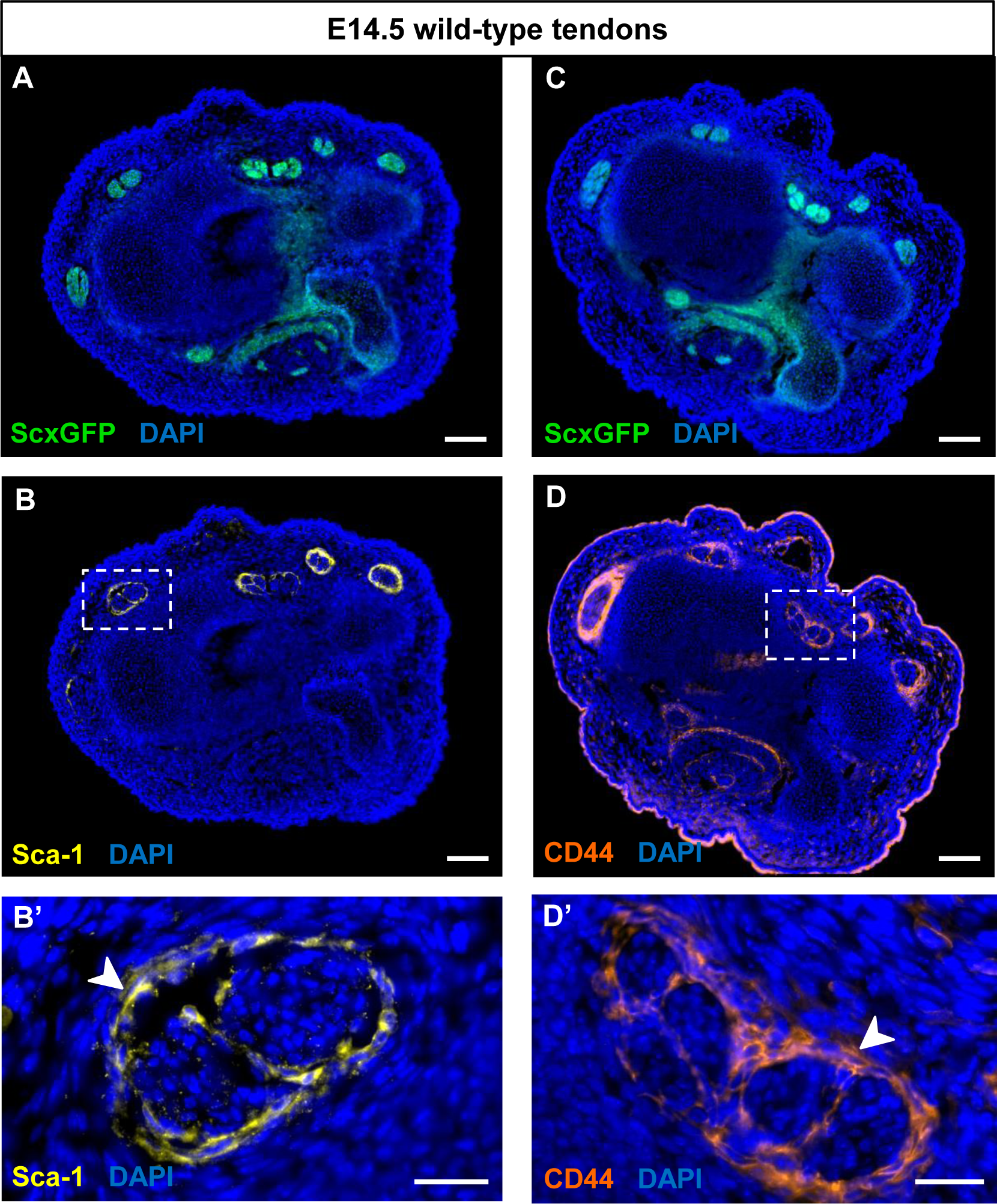
Expression of Sca-1 and CD44 during embryonic tendon development. (A-D) Immunofluorescence staining for Sca-1 and CD44 on wrist-level transverse sections from E14.5 ScxGFP-carrying wild-type embryos. Robust expression of (B) Sca-1 and (D) CD44 was detected in cells that surround the tendons at E14.5 (boxed areas), likely the precursors of the epitenon/paratenon. (B’, D’) Higher magnification views of the boxed areas in (B) and (D). The epitenon/paratenon layer is indicated by white arrowheads. Note that both markers were not expressed by the tenocytes at E14.5, the onset of tenocyte differentiation or at any other stages during embryonic tendon development (not shown). Scale bars, 100 μm (A-D) and 25 μm (B’,D’).

**Fig S5.**
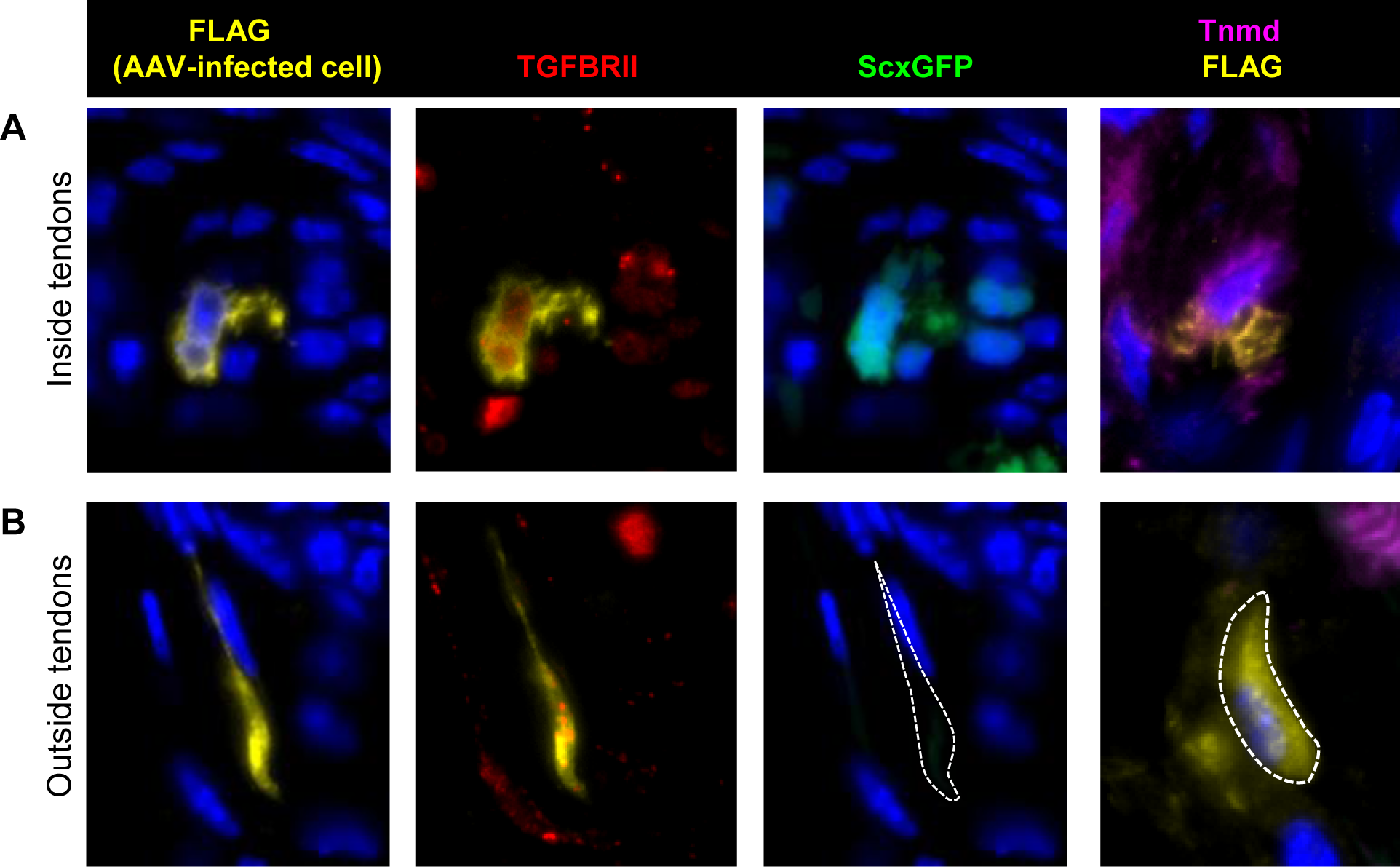
Induction of tendon markers by TGFβ signaling is context dependent. AAV1-Tgfbr2-FLAG viral infection resulted in constitutive expression Tgfbr2-FLAG expression in cells both within and outside of tendons. The virus was injected locally into P1 mutant limbs and the limbs were harvested at P7. Sections from infected limbs were stained with antibodies to FLAG (yellow) to detect infected cells, and TGFβ type II receptor (TGFBRII) to confirm the re-expression of the receptor. ScxGFP signal and tenomodulin (Tnmd) antibody staining were used to identify induction of tendon markers. (A) Infected mutant tendon cells expressed the tendon markers ScxGFP and Tnmd. (B) In cells located outside of tendons (demarcated lines), the viral infection as detected by positive FLAG and TGFBRII immunofluorescence did not result in induction of the tendon markers ScxGFP and Tnmd. Figures not to scale.

**Table S1.**
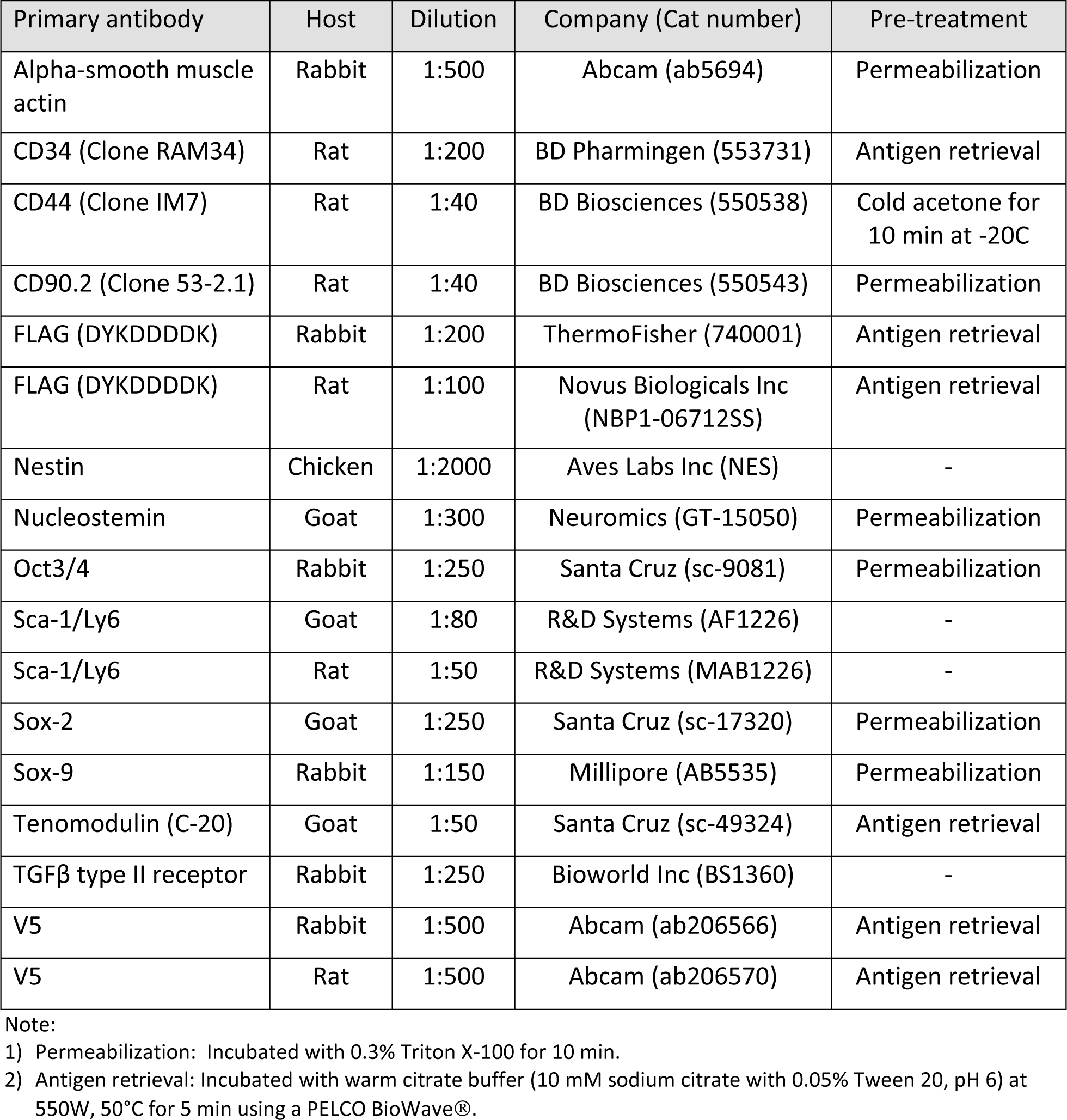
List of primary antibodies used for immunofluorescence staining.

| Primary antibody | Host | Dilution | Company (Cat number) | Pre-treatment |
| --- | --- | --- | --- | --- |
| Alpha-smooth muscle actin | Rabbit | 1:500 | Abcam (ab5694) | Permeabilization |
| CD34 (Clone RAM34) | Rat | 1:200 | BD Pharmingen (553731) | Antigen retrieval |
| CD44 (Clone IM7) | Rat | 1:40 | BD Biosciences (550538) | Cold acetone for 10 min at -20C |
| CD90.2 (Clone 53-2.1) | Rat | 1:40 | BD Biosciences (550543) | Permeabilization |
| FLAG (DYKDDDDK) | Rabbit | 1:200 | ThermoFisher (740001) | Antigen retrieval |
| FLAG (DYKDDDDK) | Rat | 1:100 | Novus Biologicals Inc (NBP1-06712SS) | Antigen retrieval |
| Nestin | Chicken | 1:2000 | Aves Labs Inc (NES) | - |
| Nucleostemin | Goat | 1:300 | Neuromics (GT-15050) | Permeabilization |
| Oct3/4 | Rabbit | 1:250 | Santa Cruz (sc-9081) | Permeabilization |
| Sca-1/Ly6 | Goat | 1:80 | R&D Systems (AF1226) | - |
| Sca-1/Ly6 | Rat | 1:50 | R&D Systems (MAB1226) | - |
| Sox-2 | Goat | 1:250 | Santa Cruz (sc-17320) | Permeabilization |
| Sox-9 | Rabbit | 1:150 | Millipore (AB5535) | Permeabilization |
| Tenomodulin (C-20) | Goat | 1:50 | Santa Cruz (sc-49324) | Antigen retrieval |
| TGFβ type II receptor | Rabbit | 1:250 | Bioworld Inc (BS1360) | - |
| V5 | Rabbit | 1:500 | Abcam (ab206566) | Antigen retrieval |
| V5 | Rat | 1:500 | Abcam (ab206570) | Antigen retrieval |

Note:
1) Permeabilization: Incubated with 0.3% Triton X-100 for 10 min.
2) Antigen retrieval: Incubated with warm citrate buffer (10 mM sodium citrate with 0.05% Tween 20, pH 6) at 550W, 50°C for 5 min using a PELCO BioWave®.

Table S2. Differentially expressed genes (2-fold change, adjusted *p*<0.05) in P7 Tgfbr2;ScxCre mutant tendon cells compared with P7 wild-type tenocytes. Note that the expression level detected for Scx also included that of ScxGFP, and therefore do not reflect the expression level of endogenous Scx.

>> See the attached excel file “Table S2-DEGs in P7 mutant cells”.

**Table S3.**
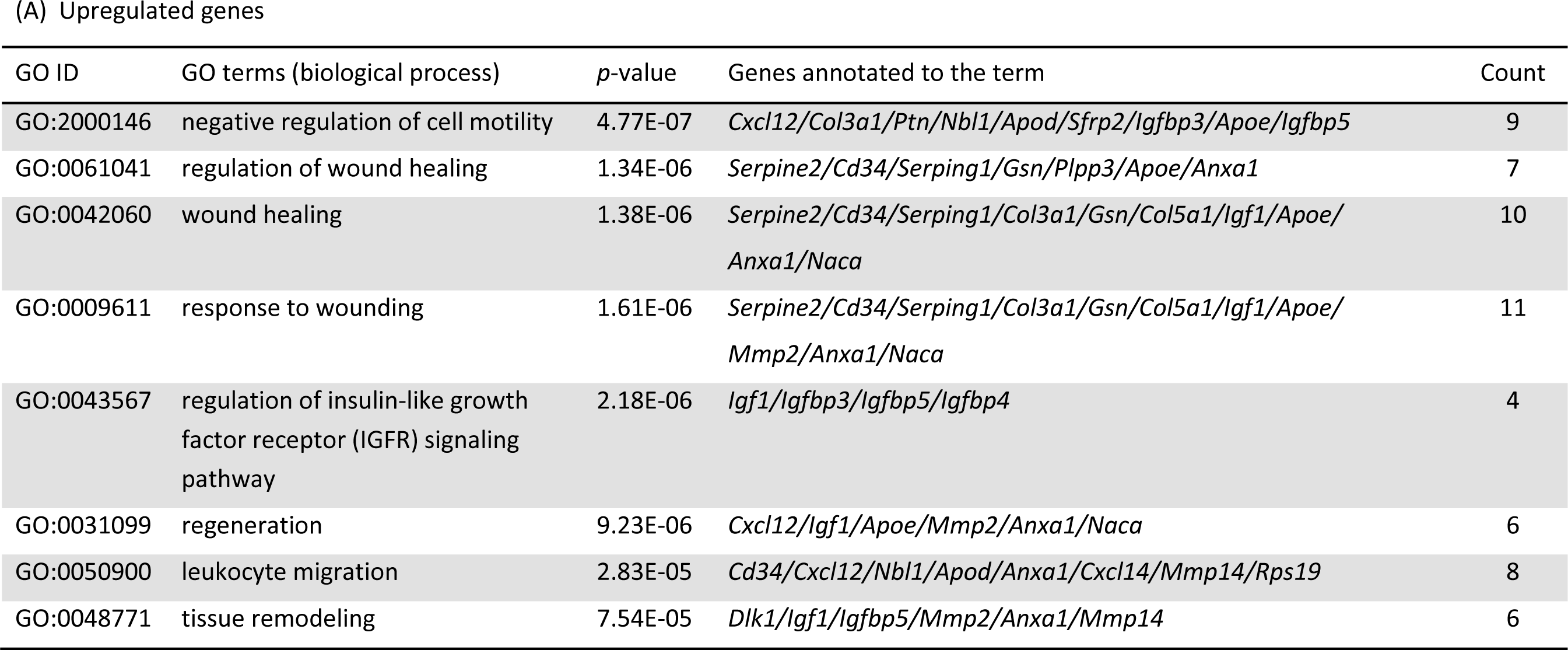
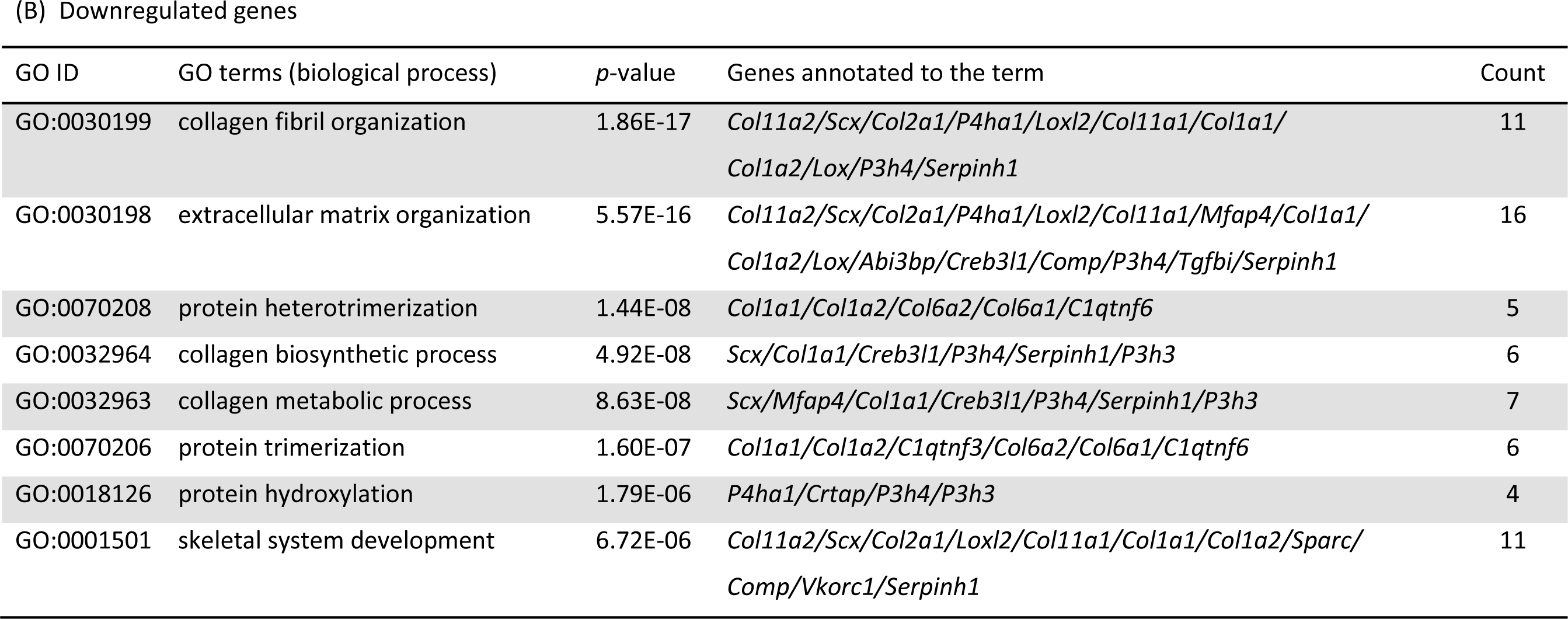
Gene Ontology (GO) term enrichment of differentially expressed genes in P7 Tgfbr2;ScxCre mutant cells compared with P7 wild-type tenocytes. A complete list of differentially expressed genes (2-fold change, *p*<0.05) used for the analysis is available in Table S2.

